# Spatially coordinated heterochromatinization of distal short tandem repeats in fragile X syndrome

**DOI:** 10.1101/2021.04.23.441217

**Authors:** Linda Zhou, Chunmin Ge, Thomas Malachowski, Ji Hun Kim, Keerthivasan Raanin Chandradoss, Chuanbin Su, Hao Wu, Alejandro Rojas, Owen Wallace, Katelyn R. Titus, Wanfeng Gong, Jennifer E. Phillips-Cremins

## Abstract

Short tandem repeat (STR) instability is causally linked to pathologic transcriptional silencing in a subset of repeat expansion disorders. In fragile X syndrome (FXS), instability of a single CGG STR tract is thought to repress *FMR1* via local DNA methylation. Here, we report the acquisition of more than ten Megabase-sized H3K9me3 domains in FXS, including a 5-8 Megabase block around *FMR1*. Distal H3K9me3 domains encompass synaptic genes with STR instability, and spatially co-localize in trans concurrently with *FMR1* CGG expansion and the dissolution of TADs. CRISPR engineering of mutation-length *FMR1* CGG to normal-length preserves heterochromatin, whereas cut-out to pre-mutation-length attenuates a subset of H3K9me3 domains. Overexpression of a pre-mutation-length CGG de-represses both *FMR1* and distal heterochromatinized genes, indicating that long-range H3K9me3-mediated silencing is exquisitely sensitive to STR length. Together, our data uncover a genome-wide surveillance mechanism by which STR tracts spatially communicate over vast distances to heterochromatinize the pathologically unstable genome in FXS.

**One-Sentence Summary:** Heterochromatinization of distal synaptic genes with repeat instability in fragile X is reversible by overexpression of a pre-mutation length CGG tract.

## Main Text

Fragile X syndrome (FXS) is the most common form of inherited intellectual disability, affecting 1 in 4,000 males and 1 in 8,000 females. The disease is made manifest early in life and presents as a range of mild to severe defects in communication skills, cognitive ability, and physical appearance, as well as hypersensitivity to stimuli, seizures, and anxiety (*1*). FXS is caused by expansion of a CGG STR tract in the 5’ untranslated (5’UTR) region of the *FMR1* gene (*2*). CGG tract length correlates with disease severity and can be stratified into <40 (normal-length), 41-55 (intermediate), 55-200 (pre-mutation), and 200+ (mutation-length) (*3-7*). Individuals with a pre-mutation length CGG tract in *FMR1* can exhibit neurodevelopmental problems in their early years, and acquire late stage neurodegeneration due to Fragile X-associated tremor/ataxia syndrome (FXTAS) (*8*). Moreover, in a process known as anticipation, mutation-length STRs grow longer as they are inherited across generations, leading to earlier onset and increased severity of FXS symptoms (*9*). These data highlight the critical role of precise STR tract lengths in a wide range of pathologic features during the onset and progression of human disease.

Increases in STR tract length correlate with pathologically altered gene expression levels in a number of repeat expansion disorders (*10*). In FXTAS, CGG expansion from normal-length to pre-mutation causes a 2-8-fold increase in *FMR1* expression, leading to pathologic nuclear inclusion bodies (*8*). By contrast, expansion from pre-mutation to mutation-length causes transcriptional inhibition of *FMR1* and consequent severe reduction in levels of the Fragile X Mental Retardation Protein (FMRP) it encodes (*11, 12*). Evidence to date suggests that transcriptional silencing occurs solely due to local DNA methylation and heterochromatinization of the *FMR1* CGG tract and its adjacent promoter (*13, 14*). Some genome-wide reports support this model by suggesting that pathologic changes to epigenetic modifications are restricted locally to *FMR1* in FXS (*15*). However, DNA demethylation by 5-aza-2’-deoxycytidine treatment or direct targeting of dCas9-Tet1 only partially reinstates *FMR1* transcription, and patient samples with longer CGG tracts are more refractory to *FMR1* de-repression (*16-18*). Moreover, recovery of FMRP levels through the use of human *FMR1* cDNA (*19*), artificial chromosomes (*20*), or viral vectors (*21-23*) cannot fully reverse FXS clinical presentations of synaptic plasticity, anxiety, seizure susceptibility, and macro-orchidism. Together, these data suggest that a subset of long-term pathologic features of FXS are made manifest independent from FMRP’s downstream effects.

We recently reported severe local misfolding of the 3D genome around the *FMR1* gene in B cells and post-mortem brain tissue from FXS patients with a 450+ CGG STR expansion (*24*), suggesting that silencing might occur via long-range mechanisms beyond local DNA methylation. Here, we investigate the extent to which 3D chromatin architecture and linear epigenetic marks are altered genome-wide as a function of a gradient of CGG STR tract lengths. We analyzed a series of human induced pluripotent stem cell lines differentiated to neural progenitor cells (iPSC-NPCs) in which the CGG STR tract is thought to expand from normal-length (5-30 CGG), pre-mutation (130-190 CGG), short mutation-length (200-300 CGG), and long mutation-length (450+ CGG Replicate 1; 450+ CGG Replicate 2) (**Fig. 1a**). To obtain precise estimates of CGG STR length, we conducted a customized assay coupling Nanopore long-read sequencing with guide RNA-directed Cas9 cutting around the transcription start site and 5’UTR of the *FMR1* gene (**Fig. 1b-e, Fig. S1, Supplementary Methods, Table S1)**. Consistent with previous reports, wild type and pre-mutation lines had on average 34 and 160 total CGG STRs, respectively, with minimal interrupting sequences (**Fig. 1b-e**). *FMR1* mRNA increased upon expansion to the pre-mutation length as previously established (**Fig. 1f**). Unexpectedly, we observed that both short- and long mutation-length FXS lines showed a similar total of 425-460 CGG triplets (**Fig. 1b**). However, the short mutation length line contained a high number of AGG interrupters, leading to shorter and less continuous CGG tracts compared to long mutation-length (green, **Fig. 1c-e**). To facilitate clarity, we refer to the three FXS lines by the sum of their top two longest continuous CGG tracts – 306, 326, and 378 CGG triplets.

**Figure 1.**
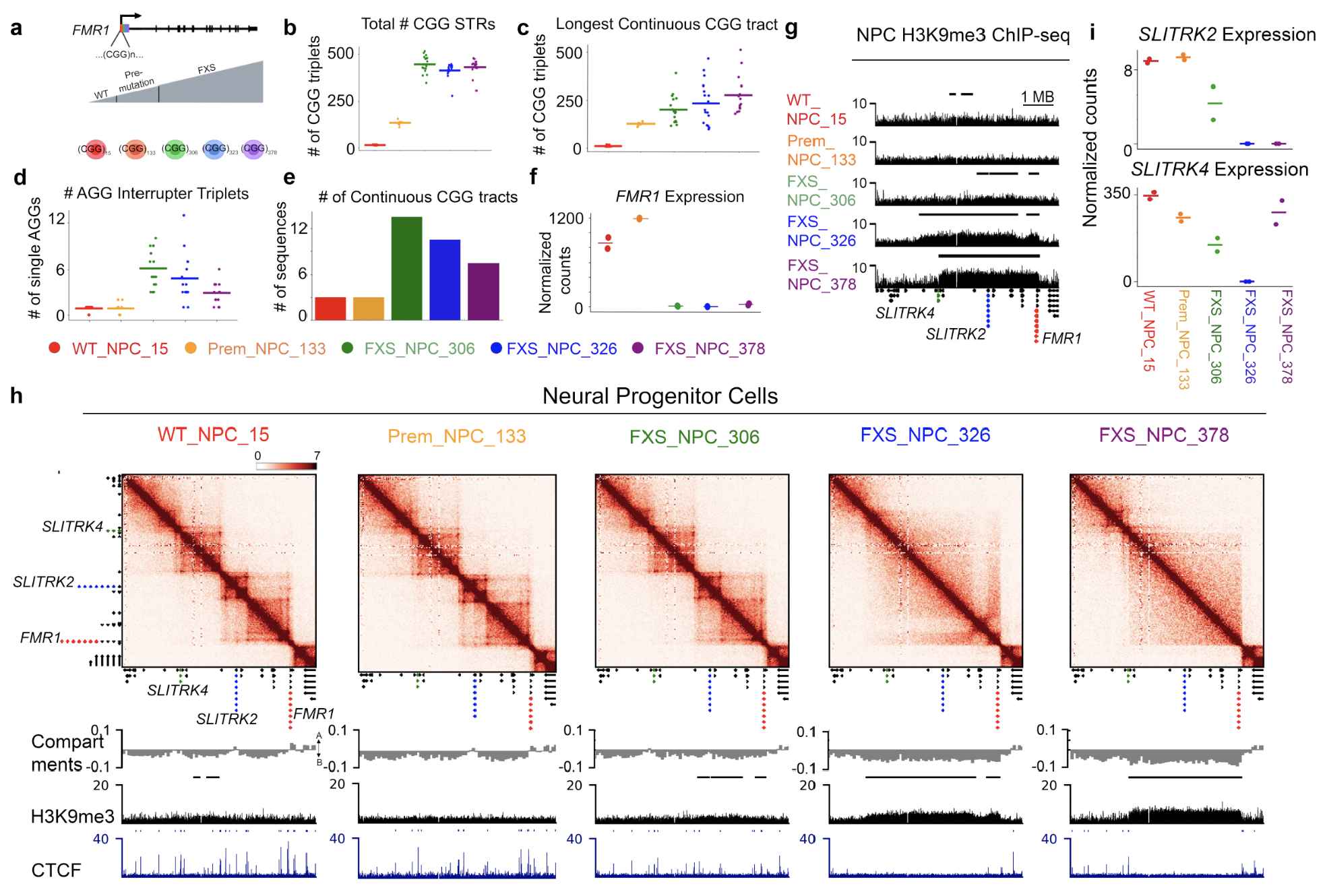
A >5 Megabase-sized domain of H3K9me3 heterochromatin spreads across the *FMR1* locus in a CGG STR length-dependent manner in fragile X syndrome. **(a)** Schematic of iPSC lines used to model FXTAS and FXS, including normal-length, pre-mutation length, short mutation-length, and long mutation-length. Colors are associated with each CGG length to identify STR sequence-dependent disease progression across all figures. **(b-e)** Nanopore long-read analysis of **(b)** total number of CGGs present, **(c)** longest continuous CGG tract, **(d)** number of AGG interrupters within CGG STR, and **(e)** total number of continuous CGG tracks within the STR in the 5’UTR of *FMR1*. **(f)** *FMR1* mRNA levels as evaluated by RNA-seq. Horizontal lines represent the central tendency (mean) between n=2 biological replicates. **(g)** H3K9me3 Chip-seq across all five lines is shown for an 8 Mb region around *FMR1*. Gene track is plotted below ChIP-seq tracks. *FMR1, SLITRK2*, and *SLITRK4* are highlighted in red, blue, and green respectively. **(h)** Hi-C data across all five lines is shown as a heatmap of counts representing interaction frequency for an 8 Mb region around *FMR1*. Compartment score, H3K9me3 Chip-seq, and CTCF Chip-seq in iPS-derived NPCs is displayed below the heatmaps for all five conditions. **(i)** *SLITRK2* and *SLITRK4* mRNA levels as evaluated by RNA-seq. Horizontal lines represent the central tendency (mean) between n=2 biological replicates.

It is well established that AGG interrupters correlate with attenuated STR instability and decreased severity of disease (*25*), therefore we hypothesized that the FXS_306 short mutation-length line with a high frequency of interrupters would have less severe pathological defects to chromatin modifications than long mutation-length lines FXS_326 and FXS_378. We observed that *FMR1* gene expression decreased significantly to the same extent in all three FXS lines (**Fig. 1f**). Concomitant with decreased *FMR1*, we observed increased DNA methylation around the transcription start site and the 5’UTR-localized CGG STR in all three FXS lines, suggesting that local levels of DNA methylation correlate strongly with the mRNA levels of *FMR1* (**Fig. S2, Table S2**). In stark contrast to local DNA methylation, we also observed CGG length-dependent acquisition of the repressive histone mark H3K9me3 (**Fig. 1g**). The gained H3K9me3 signal was not only local to *FMR1* but spread upstream over ∼3 Mb in FXS_306 and increased in strength and spread further upstream to > 5 Mb as the CGG tracts grew to 326 and 378 CGG triplets (**Fig. 1g-h, Table S3**). Thus, the spread and intensity of a large H3K9me3 repressive heterochromatin domain correlates with the length of the continuous CGG tract, whereas local DNA methylation of the *FMR1* promoter silences expression after the CGG passes short mutation-length.

We next studied the folding patterns of the 3D genome around the large acquired H3K9me3 domains (**Table S4**). In parallel with gained H3K9me3 (**Fig. 1h, Fig. S3a-b**), we observed strengthening of B compartment signal (**Fig. 1h, Fig. S3a, S3c**), loss of CTCF occupancy (**Fig. 1h, Fig. S3d**), and severe breakdown of TAD integrity (**Fig. S3a, 3e**) across the broader 5 Mb-sized H3K9me3 domain. We also observed destruction of the local subTAD boundary at *FMR1* (**Fig. S3f-h**) as previously reported in B cells and post-mortem brain tissue (*24*). Our results demonstrate that heterochromatin silencing spreads more than 5 Mb upstream of *FMR1* and correlates with severe large-scale misfolding of the 3D genome in FXS.

We noticed that the FXS H3K9me3 domain spanned two additional genes, *SLITRK2* and *SLITRK4*, encoding known neuronal cell adhesion proteins linked to synaptic plasticity (**Fig. 1g-h**). Expression of both *SLITRK2* and *SLITRK4* decreased in FXS in a manner that correlates with the spread of the H3K9me3 domain due to *FMR1* CGG expansion (**Fig. 1i**). Using our Hi-C maps, we observed that *FMR1* loops directly to *SLITRK2* and *SLITRK4* in wild type iPSC-NPCs containing a normal-length CGG STR tract (**Fig. S3i-j**). The long-range gene-gene cis interactions are abolished and *SLITRK2* and *SLITRK4* mRNA levels are decreased as H3K9me3 spreads over the locus in FXS (**Fig. S3i-n, Fig. S4**). We observed that *SLITRK2/SLITRK4* are downregulated but not fully off in the FXS_NPC_306 line, suggesting that *FMR1* silencing is governed by local DNA methylation whereas distal gene silencing is governed by general heterochromatin acquisition and 3D genome disruption (**Fig. S3i-n**). We also note that *SLITRK4* is not silenced in one of the long mutation-length samples because the H3K9me3 domain does not extend up to the promoter of the gene, further emphasizing the likely functional role for H3K9me3 in distal gene silencing in FXS (**Fig. S5**). Together, these data suggest that the acquisition of a large 5 Mb-sized H3K9me3 domain radiates outward from *FMR1* to encompass and silence additional synaptic genes as the mutation-length CGG STR further expands. FXS is characterized by clinical presentation of cognitive decline and defects in synaptic plasticity (*26*), so the direct spatial connection between *FMR1* and synaptic gene silencing is of critical importance toward understanding the onset of pathological features in the brain.

We wondered if our observations of large-scale 3D genome misfolding and heterochromatin silencing around the *FMR1* locus were specific to the NPC state. In pluripotent iPSCs, we observed the same pattern of large-scale H3K9me3 deposition gained with CGG STR expansion as in NPCs (**Fig. S6**). By contrast, in B cells our Hi-C analysis revealed that large scale genome folding disruptions did not occur upon mutation-length expansion (**Fig. S7a-b**). Importantly, the large H3K9me3 domain is pre-existing in wild type B cells with the normal-length CGG tract, but stops at the TAD boundary before *FMR1* (**Figs. S7-8**). In mutation-length FXS B cells, the pre-existing H3K9me3 domain spreads over *FMR1*, and local CTCF occupancy and TAD boundary integrity are disrupted as we have previously reported (**Fig. S7c-e**) (*24*). Thus, 3D genome folding, CTCF occupancy, and H3K9me3-based heterochromatin silencing defects are cell type-specific in FXS and most severe in cell types such as NPCs where no pre-existing H3K9me3 domain is present at the larger *FMR1* locus.

We next sought to understand if heterochromatin might be acquired on autosomes in FXS. We identified eleven additional genomic locations in which large (>1 Mb) H3K9me3 domains were acquired with low signal in FXS_306 short mutation-length and subsequently strengthened and spread in FXS_326 and FXS_378 (**Fig. 2a, Fig. S8, Tables S5-6**). The same domains were present in iPSC (**Fig. S9**). One such domain encompasses the *SHISA6* gene – a known fragile site on chromosome 17 (**Fig. 2b)**. As seen at the broader *FMR1* locus, acquisition of H3K9me3 upon mutation-length CGG expansion occurs in parallel with TAD ablation and loss of CTCF occupancy (**Fig. 2b-c**). *SHISA6* mRNA levels are decreased proportionately to the intensity of the H3K9me3 domain (**Fig. 2d**). Indeed, for all 11 distal FXS domains, we observed loss of CTCF occupancy (**Fig. 2e, Fig. S10, Table S7**), TAD boundary disruption (**Fig. 2f, Fig. S10**), and a marked reduction in gene expression (**Fig. 2g, Fig. S11**). Gene ontology analysis confirmed that genes in our FXS-specific H3K9me3 domains in iPSC-NPCs are involved in synaptic plasticity and neural cell adhesion, and such synaptic genes are not enriched in the H3K9me3 domains that are invariant across all CGG lengths (**Fig. 2i-j, Fig. S12a, Table S8**). We note that although we see both gain and loss of gene expression genome-wide in FXS (**Fig. S13**), it is only the downregulated genes in our NPC H3K9me3 domains that exhibit synaptic gene ontology (**Fig. S12a-c**). In addition to N=12 heterochromatin domains present across all FXS cell lines, we also identified 20 H3K9me3 domains specific to just one cell line (**Fig. 2a, 2j**), indicating that heterogeneity in clinical presentation in FXS patients may be due to different distributions of heterochromatinization. Together, our data reveal that large H3K9me3 domains also arise distal from the *FMR1* locus and encompass genes critically linked to the synaptic plasticity defects characteristic of FXS (*27*).

**Figure 2:**
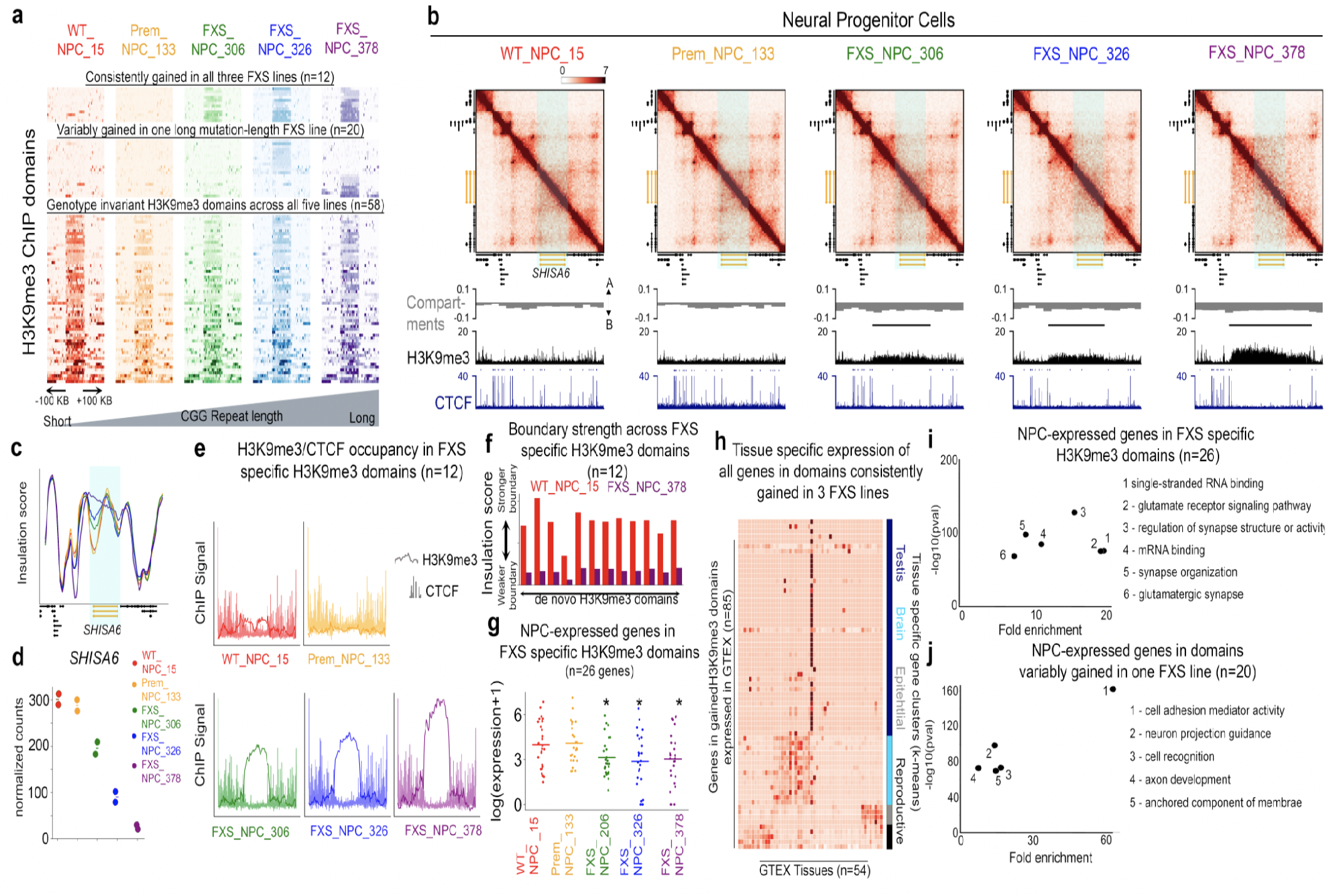
Acquisition of distal heterochromatin domains and silencing of key distal synaptic plasticity genes in fragile X syndrome. **(a)** Three classes of H3K9me3 ChIP-seq domains identified genome-wide across five iPSC-derived NPC lines. H3K9me3 domains are defined as either (i) invariant across all genotypes, (ii) gained in FXS but not consistently in all disease lines, or (iii) consistently gained in all three FXS lines. Categorization was based on presence or absence of domains identified by RSEG. **(b)** Hi-C data across all five lines is shown as a heatmap of counts representing interaction frequency for a 1.6 Mb region around one of the somatic H3K9me3 domains encompassing *SHISA6*. Compartment score, H3K9me3 ChIP-seq, and CTCF ChIP-seq in iPSC-NPCs is displayed below the heatmaps for all five conditions. Lines representing RSEG H3K9me3 domain calls are show above H3K9me3 ChIP-seq track. *SHISA6* gene is highlighted in orange. **(c)** Insulation score across Hi-C matrices in the same 1.6 Mb region as (b) for all five cell lines. Red, orange, green, blue, purple correspond to normal-length, pre-mutation, short mutation-length, long mutation-length sample 1, and long mutation-length sample 2, respectively. *SHISA6* gene is highlighted in orange. **(d)** *SHISA6* mRNA levels as evaluated by RNA-seq. Horizontal lines represent the central tendency (mean) between n=2 biological replicates. **(e)** Pooled H3K9me3 and CTCF ChIP-seq data across all n=12 consistently gained H3K9me3 domains in FXS. **(f)** Insulation score for the strongest domain boundary in each of the H3K9me3 domains consistently gained in all three FXS lines for WT_NPC_15 (red) and FXS_NPC_378 (purple) cell lines. There are 12 sets of one red and one purple bar plot, each set corresponding to one H3K9me3 domain. **(g)** mRNA levels as evaluated by RNA-seq for n=26 protein coding genes in consistently gained domains in FXS. Genes are only shown if they were expressed in at least one cell line. Each point represents expression of one gene averaged across n=2 biological replicates. * indicates p <0.05 when compared to WT_NPC_15. Pvalues were calculated using a one-tailed Mann Whitney U test. **(h)** Expression of all genes in consistently gained domains in FXS across tissues in the GTEX dataset. Genes were only shown if expression was not 0 across all tissues, resulting in n=67 genes. Genes were clustered using K-means clusters into 4 groups, and clusters were labelled based on the tissue types dominating each cluster. **(i-j)** Gene ontology (GO) analysis using WebGESTALT for **(i)** n=26 protein coding genes expressed in iPS-derived NPCs and localized to H3K9me3 domains consistently gained in FXS and **(j)** n=20 protein coding genes expressed in iPS-derived NPCs and localized to H3K9me3 domains gained in FXS but not consistently in all FXS lines as defined in panel (a).

Macro-orchidism and soft skin are unexplained clinical presentations in FXS (*28*), and expansion of the *FMR1* CGG STR also causes severe ovary defects in Fragile X-associated primary ovarian insufficiency (FXPOI) (*29*). To understand the transcriptional profile of our H3K9me3-localized genes in tissues outside the brain, we examined their expression across 54 tissues from the GTEX consortium. We observed that genes localized to FXS heterochromatin domains largely exhibit tissue-specific expression profiles, including testis, female reproductive organs, epithelium, and (consistent with our NPC results) brain (**Fig. 2h, Fig. S14**). Given that our NPC FXS-specific H3K9me3 domains are also present in iPSCs, these results suggest that many of such domains will also be present in skin and reproductive tissues and thus relevant to the silencing of genes linked to non-brain pathology. Our results bring to light a compelling hypothesis in which distal heterochromatinization and silencing of epithelial and testis genes on autosomes is a mechanism contributing to pathological features outside the brain in a broad range of clinical presentations due to *FMR1* CGG instability.

Given that the primary site of STR expansion is in *FMR1* on the X chromosome, it remains quite unusual that large distal genomic loci would be heterochromatinized in FXS. To understand how *FMR1* communicates with distal loci, we examined Hi-C trans matrices (**Supplementary Methods**). We unexpectedly observed unusually strong trans (i.e. inter-chromosomal) interactions connecting the *FMR1* locus specifically to distal H3K9me3 domains (**Fig. 3a-b, Fig. S15**). Importantly, all distal silenced H3K9me3 domains form a multi-way subnuclear hub with *FMR1* in FXS (**Fig. 3c**). We observed that the formation of trans interactions occurs concomitant with the density of H3K9me3 acquired during disease progression (**Figs. S15-20)**. Trans interactions are not present in normal-length or pre-mutation length. They initiate upon short mutation-length (FXS_306) expansion, and form full interaction strength as the H3K9me3 domains spread in long mutation-length (FXS_326, FXS_378) (**Figs. S15-20)**. Together, these data show that the genome-wide gained FXS heterochromatin domains engage directly via spatial proximity with the unstable *FMR1* locus upon mutation-length expansion of the CGG STR tract.

**Figure 3:**
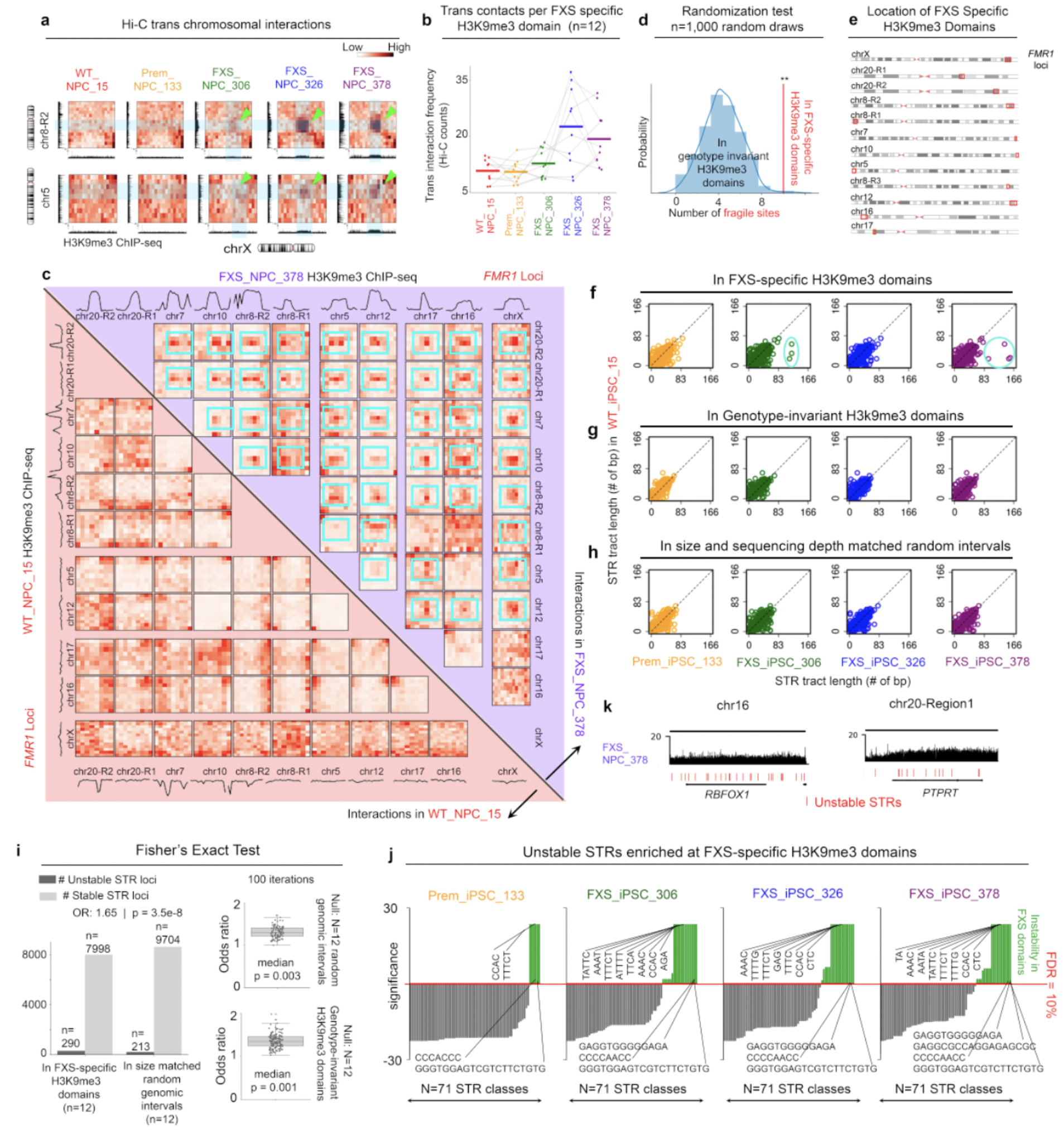
FXS heterochromatin domains form a spatial subnuclear hub of trans interactions between distal genes exhibiting short tandem repeat instability. **(a)** Hi-C inter-chromosomal interaction heatmaps binned at 1 Mb resolution between *FMR1* H3K9me3 domains and H3K9me3 domains on chromosome 8 and chromosome 5. The window for each region includes the H3K9me3 domain gained in FXS and 5 Mb of flanking genome. H3K9me3 Chip-seq is shown on the x-axis for chromosome X and for the y-axis for the distal region. Blue bars and green arrows highlight trans interactions. **(b)** Inter-chromosomal contacts between all FXS H3K9me3 domains and the *FMR1* domain on chromosome X. Each dot represents one gained H3K9me3 domain. Lines connecting the dots show the progress of that domain across 5 cell lines with increasing CGG STR length. Bar represents mean trans contacts across all domains. **(c)** Pairwise Hi-C interactions among all of the distal H3K9me3 domains gained in FXS for long mutation-length (FXS_NPC_378, upper triangle) and normal-length (WT_NPC_15, lower triangle). Domains annotated by chromosome. The window for each region includes the H3K9me3 domain gained and 3 Mb of flanking genome. H3K9me3 Chip-seq signal for all domains for both FXS_NPC_378 and WT_NPC_15 are plotted above Hi-C heatmaps. Blue boxes highlight FXS gained trans interactions. **(d)** The number of fragile sites within the H3K9me3 domains in *FXS* compared to a null distribution consisting of 1000 draws of n=12 size-matched, randomly-sampled intervals located within genotype-independent H3K9me3 domains that remain constant throughout normal-length, short mutation, premutation, and disease lines. as called by RSEG. Fragile sites were obtained from the HumCFS database. **(e)** The location of the gained H3K9me3 domain at *FMR1* and n=11 distal gained H3K9me3 domains is highlighted in a red box on a chromosome ideogram obtained from the UCSC genome browser. **(f)** The length of STR types (in bp) in FXS-specific H3K9me3 domains which differ by at least 3 units in at least 1/3 disease lines compared to WT are plotted. WT_NPC_15 length is plotted on the y-axis and remaining cell lines are plotted on the x-axis. **(g)** Same as f, for STRs in genotype-invariant H3K9me3 domains. **(h)** Same as f, but for random genomic intervals matched to FXS-specific H3K9me3 domains in size and sequencing depth. **(i)** Fisher’s Exact test comparing the number of stable vs unstable STR loci in FXS specific H3K9me3 domains vs an equal number coverage matched genome invariant H3K9me3 domains. The results of a 1000 Fisher’s exact tests, run on 1000 different sets of randomly drawn size matched genomic intervals or genome invariant H3K9me3 domains are also shown. **(j)** Results of a statistical test (see Supplemental Methods) demonstrating which of the N=71 classes of STRs which are unstable in FXS are enriched for instability in FXS-specific H3K9me3 domains vs genome invariant H3K9me3 domains. An FDR of 10% was used as a threshold for significance. The individual N=71 loci are plotted on the x-axis from least to most significant. The y-axis, significance, is - 10*log(BH-corrected p-value + pseudocount), centered so that FDR of less than 10% is positive and the remaining are negative. **(k)** H3K9me3 and unstable STRs are shown for the *RBFOX1* gene and the *PTPRT* gene.

Heterochromatinization is known to protect the repetitive genome against instability (*30*). We hypothesized that genes in FXS H3K9me3 domains would require spatially coordinated heterochromatinization because they fall in genomic locations that are highly susceptible to instability. Consistent with this idea, we first noticed that the majority of our FXS domains overlapped established human fragile sites (**Fig. 3d, Fig. S21**). We also noticed that, like *FMR1*, nearly all of the FXS-specific distal H3K9me3 domains are located at the ends of chromosomes adjacent to sub-telomeric regions (**Fig. 3e**). Using high-coverage whole genome PCR-free sequencing and the GangSTR computational method (*31*), we quantified STR tract length genome-wide across our healthy and FXS iPSC lines (**Table S9)**. We observed that a small subset of STR types are indeed expanding or contracting specifically in our FXS disease iPSC lines compared to normal-length wild type iPSCs **(Fig. 3f, Fig. S22, Supplementary Methods)**. Importantly, pathologically unstable STR tracts are highly enriched in our FXS-specific H3K9me3 domains compared to random size-matched genomic intervals or genotype-invariant H3K9me3 domains **(Fig. 3f-h, Fig. 3i, Fig. S23a, Supplementary Methods)**. Together, this work demonstrates that regions of the genome silenced in FXS are similar to *FMR1* loci in that they are at the ends of chromosomes and are enriched for pathologically unstable STRs. We posit that instability may predispose distal STR loci as targets of the same mechanisms driving H3K9me3 deposition at the larger *FMR1* locus.

To further understand why the unstable *FMR1* locus would spatially contact and coordinate heterochromatinization with our specific distal locations and not with other locations in the genome, we explored the genetic features of unstable STRs in our FXS H3K9me3 domains. We formulated a statistical test (**Supplementary Methods**) to identify specific STR tracts which are expanding at higher-frequency in our FXS-specific H3K9me3 domains compared to random size-matched genomic intervals or genotype-invariant H3K9me3 domains (**Fig. 3j, Fig. S23b**). A list of top pathologically unstable STRs enriched in FXS-specific H3K9me3 domains are provided (**Tables S11-S13)**. Distal CGG STR tracts did not noticeably expand, but this could be due to inaccurate GangSTR estimates on high CG-content tracts (**Fig. S23c**). Importantly, genes localized in FXS-specific H3K9me3 domains are significantly longer and exhibit a significantly higher density of pathologically unstable STR tracts per gene compared to the null expectation of all genes in random size-matched genomic intervals or even genotype-invariant H3K9me3 domains present across all five lines **(Fig. 3k, Fig. S23d-e)**. Together, our data inspired a working model in which STR tracts across the genome communicate with each other spatially via trans interactions as a surveillance mechanism that enables the heterochromatinization and silencing of STR tracks at risk of instability.

To understand the functional role of the *FMR1* CGG STR in heterochromatin deposition in cis and in trans, we examined the extent of H3K9me3 reversibility after shortening the CGG to pre-mutation or normal-length with CRISPR (**Fig. 4a**). In our first iPSC cohort, we cut back the *FMR1* CGG tract in the long mutation-length FXS_iPSC_378 line to the normal-length range of 4 CGG triplets as confirmed by Nanopore long-reads (**Fig. 4a-b, Figs. S24-25**). We observed that the large H3K9me3 domain spanning *SLITRK4, SLITRK2*, and *FMR1* did not notably change after CGG STR cut-back to normal-length (**Fig. 4c**). CTCF binding was not re-gained and genome folding domains remained destroyed just as in the FXS_iPSC_378 parent line (**Fig. S26**). Consistent with previous reports, the *FMR1* gene was partially de-repressed in the normal-length cut-out, however *SLITRK2* remained silenced (**Fig. 4d**). We also noticed that all distal heterochromatinized loci maintained a high level of H3K9me3 signal upon normal-length CGG cut-out (**Fig. S27**). Our data indicate that engineering the CGG STR back to normal-length range does not markedly reprogram H3K9me3 domains in cis or in trans, suggesting that pathologically silenced synaptic, epithelial, testis, and female reproductive tissue genes will not be de-repressed with an *FMR1* CGG normal-length cut-out strategy in FXS.

**Figure 4:**
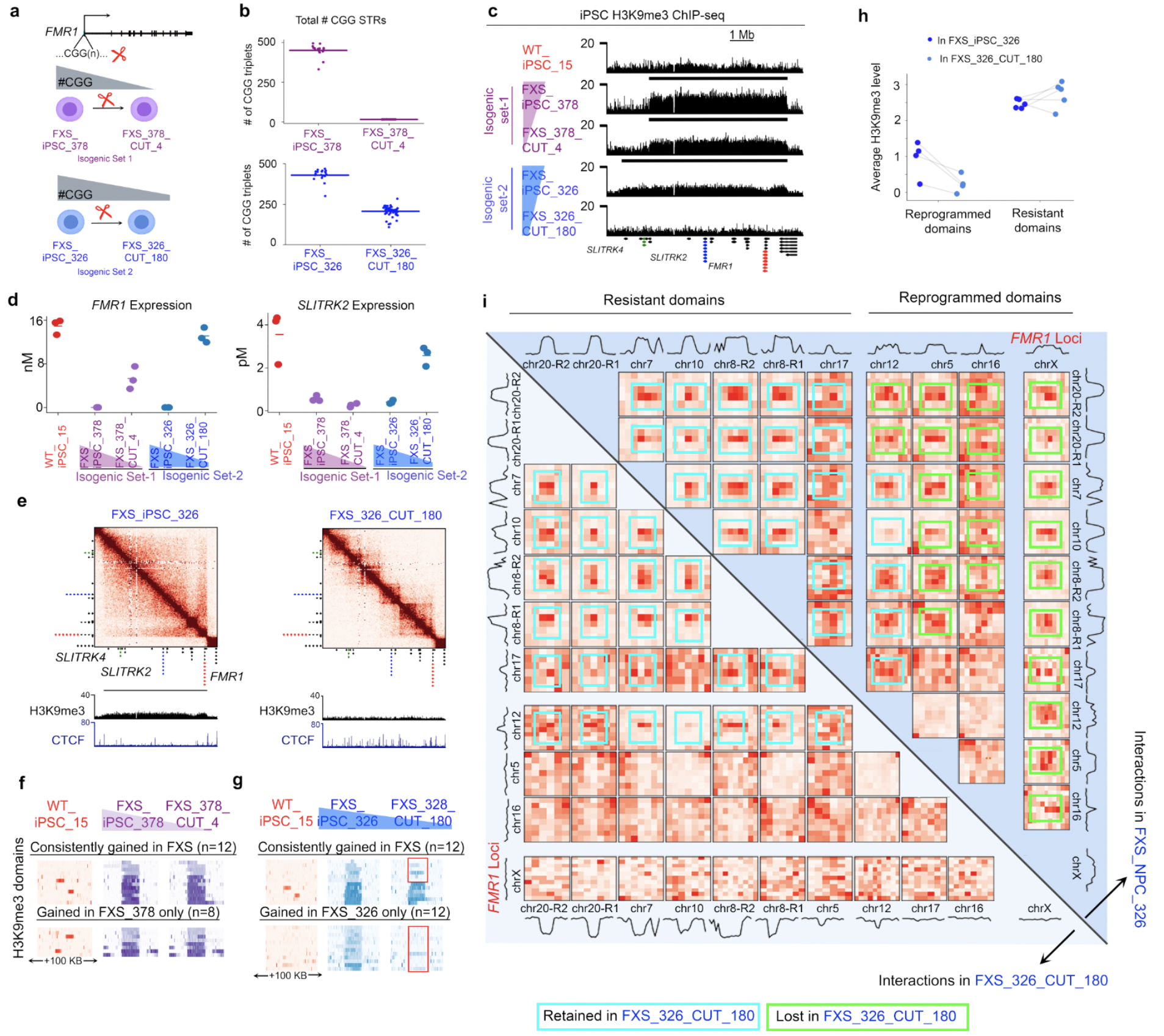
Engineering the *FMR1* CGG STR to pre-mutation length reverses a subset of distal heterochromatin domains and reprograms 3D genome misfolding in fragile X syndrome. **(a)** Schematic of CRISPR engineered STR cut-out iPSCs. Isogenic set 1 (purple) consists of long mutation-length FXS line (FXS_iPSC_376) engineered to normal-length (FXS_iPSC_376_cut_4) where CGG repeats were removed so only 4 remained. Isogenic set 2 (blue) consists of a second long mutation-length FXS line (FXS_iPSC_326) engineered to pre-mutation length (FXS_iPSC_326_cut_180) where CGG repeats were removed so only 180 remained. **(b)** Number of CGGs present in the *FMR1* 5’UTR per cell line. Each dot represents the number of CGGS in one long Nanopore read. Bar represents mean across all reads. **(c)** H3K9me3 ChIP-seq in WT iPSC and Isogenic Set 1 and Set 2 iPSC for a 8 Mb region around *FMR1*. Genes are shown below ChIP-seq tracks. *FMR1, SLITRK2*, and *SLITRK4* are highlighted in red, blue, and green respectively. **(d)** *FMR1* and *SLITRK2* mRNA levels (n=3 replicates) from qRT-PCR shown for WT, Isogenic Set 1, and Set 2 iPSCs. Each dot represents one replicate, with the horizontal line representing the mean. **(e)** Hi-C interaction frequency heatmaps in an 8 Mb region around *FMR1* for iPSC lines in isogenic set 2. H3K9me3 and CTCF ChIP-seq is displayed below the heatmaps. **(f-g)** H3K9me3 ChIP-seq signal is shown for each of n=12 heterochromatin domains consistently gained across all three FXS lines as well as heterochromatin domains present only in the FXS parent line for iPSC in **(f)** isogenic set 1 and **(g)** isogenic set 2. Each line of the heat map represents one region. Red boxes annotate reprogrammed domains that lose H3K9me3 signal upon *FMR1* CGG STR shortening. **(h)** Average H3K9me3 signal in isogenic set 2 (FXS_iPSC_326 and FXS_326_CUT_180) for each FXS H3K9me3 domain, stratified up by whether the domain was reprogrammed or resistant upon shortening of the mutation-length CGG to pre-mutation length. **(i)** Pairwise Hi-C interactions among all of the distal H3K9me3 domains gained in FXS for long mutation-length (FXS_iPSC_326, upper triangle) and pre-mutation length (FXS_180, lower triangle). Domains annotated by chromosome. The window for each region includes the H3K9me3 domain gained and 3 Mb of flanking genome. H3K9me3 ChIP-seq signal for all domains for both lines are plotted above Hi-C heatmaps. Blue boxes highlight FXS trans interactions that are resistant to reprogramming. Green boxes highlight FXS trans interactions that are reprogrammed upon CGG shortening to pre-mutation length.

In our second iPSC cohort, we cut back the *FMR1* CGG tract in the long mutation-length FXS_iPSC_326 line to a pre-mutation length of 180 CGG triplets, as confirmed by Nanopore sequencing (**Fig. 4a-b, Figs. S24-25**). We unexpectedly observed that the H3K9me3 domain encompassing *SLITRK4, SLITRK2*, and *FMR1* is fully reversible upon cut-out to premutation-length (**Fig. 4c**). Corroborating the loss of H3K9me3, CTCF occupancy was re-gained and TAD boundaries were re-instated at the broader *FMR1* locus (**Fig. 4e**). Both *SLITRK2* and *FMR1* mRNA levels were nearly fully restored upon engineering to pre-mutation length (**Fig. 4d)**. Our results suggest that the reversal of the H3k9me3 heterochromatin domain around *FMR1* might require a step back through the stage of disease acquisition involving the pre-mutation length CGG STR.

We next queried the extent to which the distal H3K9me3 domains in FXS could be reversed upon local *FMR1* CGG STR engineering. By contrast to the cut-out to normal-length range where no distal H3K9me3 signal was altered, we observed that a subset of distal H3K9me3 domains were fully reprogrammed upon only engineering of the *FMR1* CGG STR precisely to 180 CGG pre-mutation length (**Fig. 4f-g, Fig. S27)**. Distal domains with the lowest H3K9me3 density were the most susceptible to reprogramming after engineering the *FMR1* CGG STR (**Fig. 4h**). Although the majority of distal high-density H3K9me3 loci remained tethered in a trans interaction hub, the *FMR1* locus and several distal domains lost their heterochromatinization and spatially disconnected upon engineering of the mutation-length CGG at *FMR1* to pre-mutation (**Fig. 4i, Fig. S28**). Together, these results highlight the remarkable ability of the *FMR1* CGG STR to communicate spatially in trans with distal H3K9me3 domains, functionally contributing, at least in part, to the acquisition of their pathologic heterochromatinization. Importantly, reverse engineering of the *FMR1* CGG to pre-mutation length can fully reverse the H3K9me3 domain locally at *FMR1* and attenuate a subset of distal H3K9me3 domains. The persistence of heterochromatin silencing at many reprogramming resistant H3K9me3 domains in FXS highlights the importance of additional clinical interventions beyond *FMR1* CGG STR engineering, and suggests that many distal H3K9me3 domains in FXS may form through a mechanism that is independent of the *FMR1* CGG.

Finally, we sought to understand if overexpression of a pre-mutation CGG STR sequence alone, independent from its placement in the *FMR1* gene, was sufficient to attenuate local or distal FXS H3K9me3 domains. We queried gene expression and H3K9me3 after overexpressing a transgene expressing 99 CGG triplets (pre-mutation) in long mutation-length FXS iPSCs for 48 hours (**Fig. 5a**). We observed a striking de-repression of *FMR1, SLITRK2, DPPA6*, and *SHISA6*, with a much higher effect size than that observed due to CRISPR CGG engineering to pre-mutation length within the endogenous *FMR1* locus (**Fig. 5b-e, Fig. S29**). Using CUT&RUN for H3K9me3, which is amenable to assaying signal in low cell numbers, we observed complete ablation of nearly all distal H3K9me3 heterochromatin domains in FXS upon overexpression of the pre-mutation CGG STR (**Fig. 5f-h**). Altogether, these data reveal that both local and distal heterochromatin domain acquisition in FXS can be fully reversed by ectopic expression of a pre-mutation length CGG STR, suggesting that the spatial subnuclear hub of fragile repetitive regions in FXS is driven by a CGG-mediated DNA or RNA mechanism that transcends *FMR1*.

**Figure 5:**
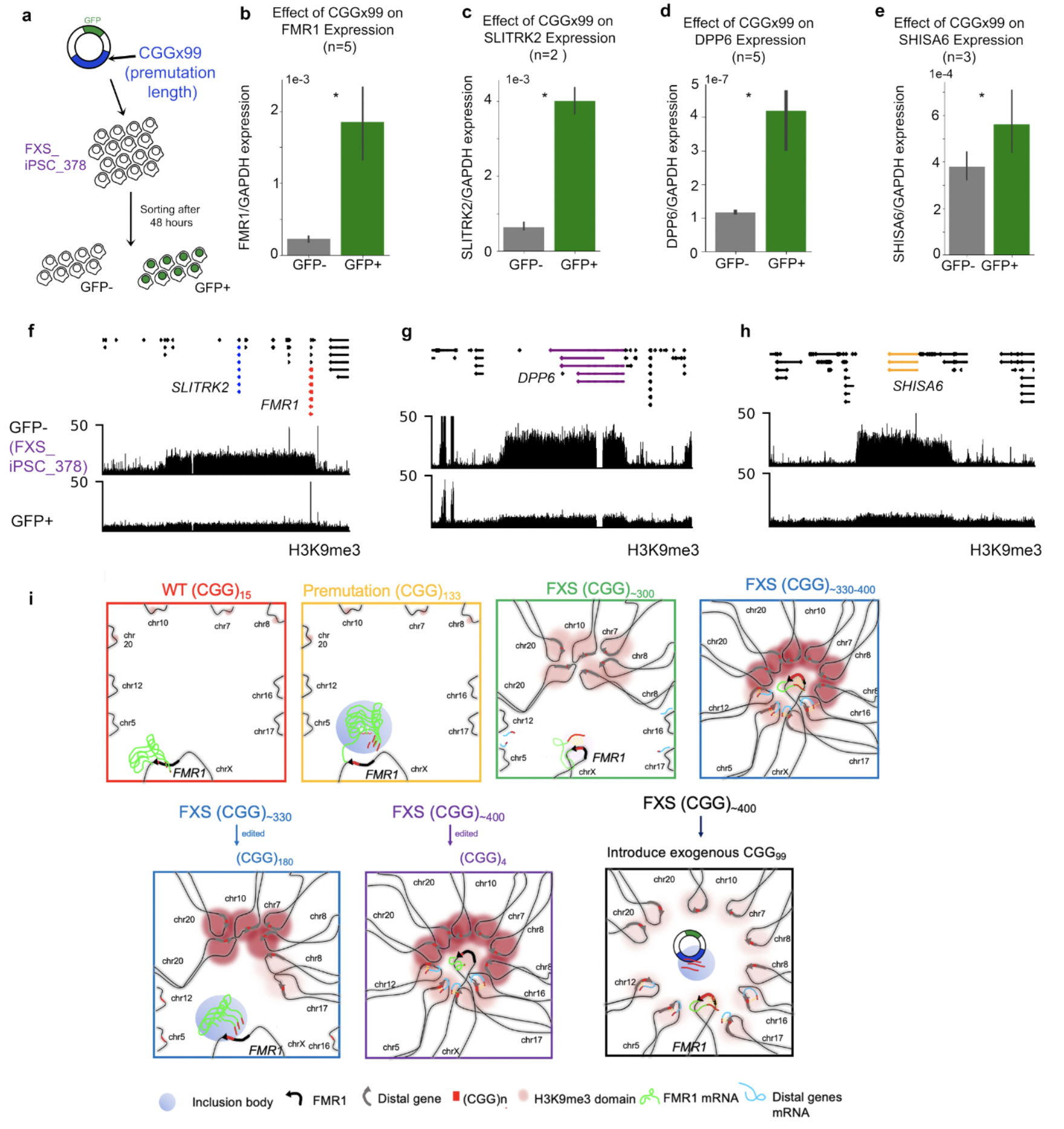
Overexpression of a pre-mutation length CGG STR tract de-represses pathologically silenced expression and attenuates FXS H3K9me3 domains. Schematic showing experimental workflow. **(b-e)** mRNA levels as assessed by qRT-PCR for **(b)** *FMR1*, **(c)** *SLITRK2*, **(d)** *DPP6*, and **(e)** *SHISA6* in long mutation-length FXS iPSCs which either did receive (GFP+) or did not receive (GFP-) the CGGx99 plasmid. Error bars represent the standard error of the mean for the indicated number of technical replicates. **(f-h)** H3K9me3 CUT&RUN in either the GFP- or GFP+ cells at the gained H3K9me3 domains at **(f)** *FMR1*, **(g)** *DPP6*, and **(h)** *SHISA6*. **(i)** Schematic model. We find that numerous heterochromatin domains interact via long-range trans interactions to form an inter-chromosomal subnuclear hub with the *FMR1* locus in fragile X syndrome. When CGG STRs are normal-length, *FMR1* and other chromosomes do not cluster and do not interact in trans. As the CGG tract expands to pre-mutation length, *FMR1* mRNA levels increase. When the CGG STR tract expands to short mutation-length, *FMR1* expression drastically decreases. Distal fragile sites acquire large H3K9me3 domains that cluster together spatially in trans. When CGG repeats expand to long mutation-length (450), *FMR1* mRNA levels are fully repressed, and the distal heterochromatin domains gain H3K9me3 signal intensity. Upon cutout from long mutation-length to pre-mutation, a subset of distal domains loses H3K9me3 signal and the long-range trans interactions with FMR1 are abolished. By contrast, cutout of long mutation-length to normal-length CGG triplets does not reverse heterochromatin domains, trans interactions remain connected, and genes remain repressed. Finally, the role for the pre-mutation length CGG tract is made evident upon introduction of an exogenous 99 CGG triplet STR transgene to FXS iPSCs. The presence of transcribed CGG plasmid leads to reduction of heterochromatin across all gained H3K9me3 domains and reactivates *FMR1* and distal gene expression, suggesting that long-range 3D Epigenome miswiring in FXS is driven by the DNA or RNA CGG STR sequence.

Altogether, our data support a model in which we find pervasive long-range transcriptional silencing in FXS via the acquisition of a physically connected subnuclear hub of more than ten Megabase-sized domains of the repressive histone modification H3K9me3. Such domains acquire low levels of H3K9me3 signal in the transition from pre-mutation to short mutation-length and increase in severity and spread of H3K9me3 density as the *FMR1* CGG STR expands to long mutation-length (**Fig. 5i**). Consistent with previous reports, we see that local DNA methylation of the *FMR1* gene correlates with its degree of silencing. By contrast, a large cohort of genes distal from *FMR1* are repressed in FXS in a manner commensurate with the severity of H3K9me3 deposition. It has long been thought that global gene expression disruption in FXS is due to the downstream effects of FMRP loss, however here we see that the CGG STR expansion in *FMR1* activates a genome-wide surveillance system to deposit large H3K9me3 domains to directly silence STR-rich genes localized at the ends of distal chromosomes. The FXS pathologic heterochromatin domains encompass and silence genes critical for synaptic plasticity, testis development, female reproductive system functioning, and epithelial tissue structure, which are directly related to the clinical presentations in FXS. Our results suggest that pharmacological and RNA-based interventions to reverse distal H3K9me3 silencing may provide tangible therapeutic benefits to FXS patients if genome stability can be maintained.

It is difficult to envision how a CGG STR expansion event in *FMR1* could coordinate heterochromatinization on 10 other chromosomes. Here, we see evidence of a physically linked subnuclear hub of inter-chromosomal interactions among known human fragile sites and long genes with high density of unstable short tandem repeat tracts in FXS. We hypothesize that critical areas of the genome communicate to coordinate silencing when instability events are detected. CRISPR engineering of the long mutation-length CGG tract to pre-mutation length provides evidence that at least a subset of distal domains is heterochromatinized and spatially connected in a manner that depends on the length of the *FMR1* CGG STR. We hypothesize that the DNA sequence or RNA encoded by the pre-mutation CGG STR tract will functionally contribute to the reversal of FXS heterochromatinization, as we demonstrate that overexpression of a generic pre-mutation-length CGG STR transgene results in complete attenuation of all distal H3K9me3 domains and full de-repression of distal genes. It is noteworthy that CRISPR shortening of the mutation length CGG STR to normal-length only slightly de-represses *FMR1* and has no noticeable effect on distal heterochromatin domains. Other studies showing stronger *FMR1* de-repression upon local CGG cut-out to normal-length may have started with a shorter mutation-length tract with lower density of H3K9me3 signal more amenable to reprogramming (*32, 33*). Our results suggest that genetically engineering approaches relying only on cut-out of the *FMR1* CGG may not reverse the silencing of key genes contributing to persistent pathology in FXS patients. Full attenuation of pathologic features across multiple tissues may require combination therapies coupling pharmacological intervention targeting epigenetic writers and erasers, as well as STR engineering. Altogether, our work uncovers a pervasive genome-wide surveillance mechanism by which fragile sites and STR tracts in the human genome spatially communicate over vast distances to heterochromatinize and silence the unstable genome.

## Acknowledgments

We thank members of the Cremins lab for helpful discussions, in particular Kenneth Pham and Zoltan Simandi for critical feedback and Michael Guo for assistance with GangSTR. Jennifer E. Phillips-Cremins is a New York Stem Cell Foundation – Robertson Investigator and an Alfred P. Sloan Foundation Fellow. Linda Zhou is a Blavatnik Family Fellow.

