## Supplemental Figures and Methods for "Spatially coordinated heterochromatinization of distal short tandem repeats in fragile X syndrome"

### Materials and Methods

#### Cell Culture

##### *B-lymphocytes*

Patient-derived B-lymphocytes were cultured as previously described (*1*). In brief, cells were grown in suspension in RPMI 1640 media (Sigma, R8758) supplemented with 2 mM glutamine, 15 % (v/v) Fetal Bovine Serum, and 1 % (v/v) penicillin-streptomycin (Thermo Fisher, 15140122) at 37 °C and 5 % $CO_2$. Cells were passaged every 2-4 days, when they reached a density of approximately 5 x $10^5$ cells/mL. All cell lines were male.

##### *Induced Pluripotent Stem Cells (iPSC)*

All human IPSC were obtained from Fulcrum Therapeutics (MA, USA). Cells were cultured in mTeSR Plus (STEMCELL Technology, 05825) supplemented with 1 % (v/v) penicillin-streptomycin (Thermo Fisher, 15140122) at 37 °C and 5 % $CO_2$ on Matrigel-coated plates. Cells were passaged by incubating in 5 mL of Versene Solution (Thermo Fisher, 15040066) at 37 °C for 3 min, after which Versene was inactivated by mixing with 10 mL of full growth media. Cells were passaged every 2-7 days. All iPSC culture plates were coated with 1.2 % (v/v) Matrigel hESC-Qualified Matrix (Corning, 354277) in DMEM/F-12 (Thermo Fisher, 11320033) for at least 1 hr at 37 °C. All cell lines were male.

##### *Neural Progenitor Cell (NPC) differentiation*

Human IPSC were differentiated into NPCs using a previously established protocol (*2*). Briefly, undifferentiated cells were maintained in mTeSR Plus (STEMCELL Technology, 05825) on Matrigel-coated plates. They were seeded onto fresh Matrigel plates in NPC media at a density of

16,000 cells/cm$^2$. NPC media was changed every day, and cells were harvested at the end of day 8. The NPC differentiation medium consists of DMEM/F-12 (Thermo Fisher, 11320033) with 5 μg/mL insulin, 64 μg/mL L-ascorbic acid, 14 ng/mL sodium selenite, 10.7 ug/mL Holo-transferrin, 543 μg/mL sodium bicarbonate, 10 μM SB431542, and 100 ng/mL Noggin.

##### *FMR1 CGG cut-out isogenic iPSC engineering*

The FXS_378_CUT_4 isogenic iPSC cell line (CGG cut-out from FXS_iPSC_378) was generated using CRISPR/Cas9 mediated targeted CGG deletion as described (*3*). To generate FXS_326_CUT_180, the FXS iPSC_326 parental line was cultured in Geltrex coated T75 flask. The day before electroporation, cells were fed with fresh Stemflex™ medium with 1X RevitaCell supplements. Cells were dissociated with 5 mL Accutase™ cell dissociation reagent (STEMCELL technology, 07920). After washing once with PBS, cells were resuspended in Resuspension Buffer R (Neon™ Transfection System 100 μL Kit, Invitrogen, 10431915) to a final cell density ~ $10^8$/mL. Dissociated iPSCs were then incubated with 60 ug of a plasmid containing Cas9 and gRNA targeted to the 5' end of exon 1 in *FMR1* (sequence: 5'- TGACGGAGGCGCCGCTGCCA-3'). The resulting solution was electroporated with the following parameters: pulse voltage 1,100 V, pulse width 30 ms, and pulse number 1 with cell density at 1 x $10^8$ cells/mL.

After electroporation, cells were plated into a Geltrex-coated T75 flask using Stemflex™ medium with 1 X RevitaCell supplements. On day 3 post-electroporation, cells were dissociated with Accutase for fluorescence activated cell sorting to enrich the GFP+ population and re-plated onto Geltrex-coated 10 cm Petri dishes at ~ 5k cells/plate. 1 X RevitaCell was supplemented in the Stemflex™ medium to enhance the cell viability. iPSC cell colonies were hand-picked and expanded in Stemflex™ medium from 96 wells to 12 wells and further expanded for cryopreservation.

#### Genomics assays

##### *Cell fixation*

Cells were fixed as previously described for all downstream ChIP-seq, Hi-C, and 5C assays (*1, 4-9*). Cell lines were fixed in 1 % (v/v) formaldehyde for 10 min at room temperature in either RPMI 1640 (Sigma, R8758) or in DMEM/F-12 (Thermo Fisher, 11320033) for B-lymphocytes or iPSCs/NPCs, respectively. The complete fixation media was 50 mM HEPES-KOH (pH 7.5), 100 mM NaCl, 1 mM EDTA, 0.5 mM EGTA, 11 % formaldehyde. Fixation was quenched in 125 mM glycine for 5 min at room temperature, following by 15 min at 4 °C. Crosslinked cells were washed in pre-chilled PBS before flash frozen and stored at −80 °C.

##### *Chromatin Immunoprecipitation (ChIP-seq)*

ChIP-seq was performed as previously described with minor modification (*1, 4-9*). Briefly, crosslinked cell pellets (consisting of 10 million cells for CTCF ChIP-seq or 3 million cells for H3K9me3 ChIP-seq), were lysed in cell lysis buffer (10 mM Tris pH 8.0, 10 mM NaCl, 0.2 % NP-40/Igepal, Protease Inhibitor, PMSF) on ice for 10 min. The suspension was then homogenized with pestle A 30 times. The nuclei were pelleted from the initial lysate at 2,500 g at 4 °C, and the resulting nuclei were further lysed in 500 µL of nuclear lysis buffer (50 mM Tris pH 8.0, 10 mM EDTA, 1 % SDS, Protease Inhibitor, PMSF) and incubated on ice for 20 min. Lysed nuclei were then sonicated by adding 300 µP IP Dilution Buffer (20 mM Tris pH 8.0, 2 mM EDTA, 150 mM NaCl, 1 % Triton X-100, 0.01 % SDS, Protease Inhibitor, PMSF) and transferred to sonication tubes. Samples were sonicated using a QSonica Q800R2 sonicator for 1 hr set at 100% amplitude, with the pulse set to 30 seconds on and 30 seconds off. The sonicated lysate was then pelleted at 14,000 RPM in 4 °C, and the supernatant was transferred to a reaction consisting of 3.7 mL IP Dilution Buffer, 500 μL Nuclear Lysis Buffer, 175 μL of a 1:1 ratio of ProteinA:ProteinG bead slurry (Thermofisher, 15918014 and 15920010, respectively), and 50 μg of rabbit IgG for preclearing. The preclearing reactions were rotated at 4 °C for 2 hrs. 200 uL of the pre-clearing reactions was saved as the "input" control. The remaining solution was added to an immunoprecipitation reaction consisting of 1 mL cold PBS, 20 μL Protein A, 20 μL Protein G, and 1 uL/million cells of either CTCF or H3K9me3 antibody and rotated overnight at 4 °C.

Immunoprecipitation reactions were prepared one day before cell lysis and rotated overnight at 4 °C. The next day, IP reactions were pelleted, and the supernatant was discarded. The remaining pellet was washed once with IP Wash Buffer 1 (20 mM Tris pH 8, 2 mM EDTA, 50 mM NaCl, 1 % Triton X-100, 0.1 % SDS), twice with High Salt Buffer (20 mM Tris pH 8, 2 mM EDTA, 500 mM NaCl, 1 % Triton X-100, 0.01 % SDS), once with IP Wash Buffer 2 (10 mM Tris pH 8, 1 mM EDTA, 0.25 M LiCl, 1 % NP-40/Igepal, 1 % sodium deoxycholate), and twice with TE buffer (10 mM Tris pH 8, 1 mM EDTA pH 8). The IP DNA was eluted from the washed beads in Elution buffer (100 mM $NaHCO_3$, 1 % SDS, prepared fresh) by resuspending and then spinning at 7,500 RPM. RNA was degraded with 2 uL RNase A (Sigma, 10109142001) and incubated at 65 °C for 1 hr. To degrade residual DNA, 3 uL proteinase K (NEB P8107S) was added, and all samples were incubated overnight at 65 °C. DNA was extracted using phenol:chloroform and ethanol precipitation methods. Antibodies used in this study were: CTCF (Millipore 07-729), H3K9me3 (Abcam ab8898), H3K27ac (Abcam ab4729), H3K27me3 (Millipore 07-449), IgG (Sigma I8140).

### *Hi-C*

We prepared Hi-C libraries using the Arima Genomics Hi-C kit (Arima Genomics, A510008) according to the manufacturer's protocol. Briefly, we enzymatically digested genomic DNA within nuclei of crosslinked cell pellets and created biotinylated ligation junctions between the digested ends at proximity. Then we extracted DNA and sheared to an average size of ~400 bp using a Covaris S220 sonicator at 140 W peak incident power, 10 % duty factor, and 200 cycles per burst for 55 seconds. We further size selected the sheared DNA to 200-600 bp using AgenCourt Ampure XP beads (Beckman Coulter, A63881) according to the manufacturer's protocols. Biotin-tagged ligation junctions were pulled down using streptavidin breads from the Arima Hi-C kit according to the manufacturer's protocol. Streptavidin beads containing Hi-C libraries were stored at -20 °C for no more than 3 days before Illumina sequencing library preparation was performed.

##### *Chromosome-Conformation-Capture-Carbon-Copy (5C)*

<u>In Situ 3C</u>

3C libraries were prepared as previously described (*1, 4-9*). In brief, crosslinked cell pellets were lysed in cell lysis buffer (10 mM Tris pH8.0, 10 mM NaCl, 0.2 % (v/v) NP-40) and supplemented with 17 % (v/v) Protease inhibitor cocktail (Sigma, P8340) on ice for 15 min. Nuclei were isolated by centrifuging cell lysate at 2,500 g for 5 min at 4 °C. Pellets were washed once in cell lysis buffer and permeabilized in 0.5 % (w/v) SDS at 65 °C for 10 min. SDS was quenched in 6.6 % (v/v) TritonX-100 at 37 °C for 15 min. To create 3C ligation junctions, chromatin was digested using 100 U of HindIII in NEBuffer 2 (NEB, B7002S) at 37 °C overnight and inactivated at 62 °C for 30 min after overnight incubation. Digested ends at proximity were ligated using 1,000 U T4 DNA ligase (NEB, M0202S) in 1 X T4 DNA ligase buffer supplemented with 0.83 % (v/v) TritonX-100 and 0.1 mg/mL BSA at 16 °C for 2 hrs. The reaction was spun down at 2,500 g for 5 min, the supernatant was discarded, and the pellet was resuspended in nuclear lysis buffer (10 mM Tris-Hcl pH 8.0, 0.5 M NaCl, 1.0 % SDS). Crosslinks were reversed with the addition of 25 µL of 20 mg/mL Proteinase K (NEB, P8107) and incubated at 65 °C for 4 hrs. An additional 25 uL of Proteinase K was then added and incubated at 65 °C overnight. RNA was degraded in 0.3 mg/mL of RNase A at 37 °C for 30 min. DNA was extracted with 350 µL phenol:chloroform and precipitated with sodium acetate and ethanol. Excess salt was removed using Amicon Ultra centrifugal filter unit (Millipore, MFC5030BKS).

5C

5C libraries were prepared as previously described (*1, 4-9*). In brief, we used previously designed double alternating 5C primers to a 6.4 Mb-sized region around the *FMR1* locus (*1*). 1 fmole of 5C primers were denatured at 95 °C for 5 min and then annealed to 600 ng of 3C template in 1 X NEBuffer 4 (NEB, B7004S) at 55 °C for 16 hrs. Annealed 5C primers were ligated by 10 U of Taq Ligase (NEB, M0208L) at 55 °C for 1 hr. Ligase was inactivated at 75 °C for 10 min, followed by PCR amplification in PCR mix (5 µL 5 X HF buffer, 0.2 µL 25 mM dNTP, 1.5 µL 80 µM emulsion forward primers, 1.5 µL 80 µM emulsion phosphorylated reverse primers, 0.25 µL Phusion polymerase (NEB, M0530L), 10.55 µL nuclease-free water) in 3 stages: 1 cycle 95 °C for 5 min; 30 cycles - 98 °C for 10 seconds, 62 °C for 30 seconds, 72 °C for 30 seconds; 1 cycle 72 °C for 10 min; and 4 °C hold. 5C libraries were then prepared for sequencing with Illumina sequencing library preparation.

***Total RNA-seq***

We isolated total RNA from NPCs and IPSCs using the mirVana miRNA Isolation Kit (Thermo Fisher, AM1560) according to the manufacturer's protocol. We used 100 ng of isolated RNA for RNA-seq library preparation using TruSeq Stranded Total RNA Library Prep Gold (Illumina, 20020598) according to the manufacturer's instruction. In brief, we removed rRNA from the input RNA, followed by double stranded cDNA preparation using 0.8 U of SuperScript II RT (Thermo Fisher, 4376600) and A-tailing end repair. We ligated cDNA to TruSeq RNA Single Indexes Set A (Illumina, 20020492) to enable multiplex sequencing and performed one round of size selection (selecting for 300 bp) and bead clean-up. 42.5 uL of sample was purified with 42 uL of Agencourt AMPure XP beads (Beckman Coulter, A63881), and 50 uL of sample was cleaned with 50 uL of Agencourt AMPure XP beads (Beckman Coulter, A63881). We amplified the purified samples using 15 PCR cycles, and samples were further purified using Agencourt AMPure XP beads (Beckman Coulter, A63881). We assessed library quality and quantities using the Agilent DNA 1000 reagent kit (Agilent, 5067–1504) on the Agilent Bioanalyzer 2100 (Agilent, 5067–4626) and using the Qubit high sensitivity RNA assay kit (Thermo Fisher, Q32852) on the Qubit Fluorometer, respectively, before sequencing on NextSeq500 (Illumina).

***High throughput DNA sequencing – library preparation***

ChIP-seq and 5C libraries were prepared for sequencing using the NEBNext Ultra II DNA Library Prep Kit (NEB #7103) according to the manufacturer's protocol. For ChIP-seq and 5C, size selection of adaptor-ligated libraries was performed using AgenCourt Ampure XP beads (Beckman Coulter, A63881) according to the manufacturer's protocol. For 5C, size selection targeted ~230 bp fragment size and libraries were amplified using 5 PCR cycles. For ChIP-seq, size selection targeted <1 kb fragment size and libraries were amplified using 11 PCR cycles. Input amounts for library preparation using the NEBNext Ultra II DNA Library Prep Kit were 1 ng of purified ChIP-seq libraries and 100 ng of purified 5C libraries.

Hi-C libraries were prepped for sequencing by washing adaptor-ligated Hi-C libraries on streptavidin beads twice in 150 uL of wash buffer at 55 °C and once in 100 mL of elution buffer at room temperature using an Arima Hi-C kit (Arima Genomics, A510008). DNA was eluted from streptavidin beads by boiling at 98 °C for 10 min in 15 uL elution buffer. Subsequently, the libraries were amplified using NEBNext Ultra II DNA Library Prep Kit for Illumina (NEB, E7645S) with 8 PCR cycles according to the manufacturer's protocol.

RNA-seq libraries were prepared for sequencing using the TruSeq Stranded Total RNA Library Prep Gold (Illumina, 20020598) according to the manufacture's protocol.

##### *Sequencing*

Prior to sequencing on an Illumina NextSeq 500, library quality and size distribution were analyzed with Agilent Bioanalyzer High Sensitivity DNA Analysis Kits (Agilent, 5067-4626) and quantified using Kapa Library Quantification Kit (KAPA biosysytem, KK4835). ChIP-seq libraries were sequenced with 75 bp single end reads. 5C and Hi-C libraries were sequenced with reading length 37 bp paired end reads. RNA-seq libraries were sequenced with 75 bp paired-end reads.

##### *Gene expression quantification using qRT-PCR*

Genes of interest were quantified as previously described (*1*). Briefly, RNA isolation was performed on IPSCs and NPCs by harvesting cells, snap freezing them in liquid nitrogen, and storing at -80 °C until RNA extraction. 1 x $10^6$ frozen cells were thawed on ice, and total RNA was extracted using mirVana™ miRNA Isolation Kit (Thermo Fisher, AM1560) according to the manufacturer's protocol. RNA was converted into cDNA for each sample using the SuperScript® First-Strand Synthesis System for RT-PCR (Thermo Fisher, 11904018) according to the manufacturer's instruction. 100 ng of RNA was used as input for each sample, and RNA was quantified using the Qubit RNA HS assay (Thermo Fisher, Q32852).

To perform qRT-PCR reactions, 2 uL of cDNA was mixed with 10 uM forward and 10 mM reverse primers for a final concentration of 400 nM, in 1X Power SYBR Green PCR Master Mix (Thermo Fisher, 4368706), and the reaction was completed on the Applied Biosystems StepOnePlus Real-Time PCR System (Thermo Fisher, 4376600) according to the manufacturer's instructions. qPCR conditions were 95 °C for 10 min, followed by 40 cycles of 95 °C for 15 seconds and 65 °C for 45 seconds. Primer pair specificity was validated with single-peak melting curves at the end of PCR cycles. Standards were created with serial dilutions of 200–0.0002 pM cDNA. The resulting CT values were used to generate a standard curve and compute the concentration of mRNA transcripts per condition using 100 ng of RNA in the cDNA reaction.

##### ***Nanopore long-read sequencing of CGG short tandem repeat tract in FMR1***

High-molecular-weight DNA preparation

We conducted long-read sequencing on the *FMR1* locus using a protocol modified from previous work (*10, 11*). Briefly, 1 x $10^7$ iPSCs were resuspended in 100 µL of 1 X PBS. Cells were lysed by adding 10 mL of TLB solution composed of 10 mM Tris-Cl (pH 8), 25 mM EDTA (pH 8), 0.5 % SDS (w/v), and 20 µg/mL RNase A (Sigma) for 1 hr at 37 °C. Then, proteins were digested at 50 °C for 3 hrs using 50 µL of Proteinase K (BIO-37084). The viscous solution was transferred into a 50 mL Falcon tube containing 5 g of phase-lock gel, and 10 mL of ultrapure

Phenol/Chloroform/Isoamyl Alcohol (Fisher) was added. Samples were mixed on a rotator at 40 RPM for 10 min. Phase separation was performed by centrifugation at 2,800 g for 10 min. The aqueous phase was then carefully poured into a fresh 50 mL Falcon tube containing 5 g of phase-lock gel, followed by a second phase separation using 10 mL of ultrapure Phenol/Chloroform/Isoamyl Alcohol. Samples were mixed and centrifuged as described above. The aqueous phase was poured into a fresh 50 mL Falcon tube, and the genomic DNA was precipitated using 4 mL of 5 M ammonium acetate together with 30 mL of ice-cold ethanol (100%) and gently inverted ten to twenty times. Precipitated DNA was centrifuged at 12,000 g for 5 min and washed with 70 % ethanol twice. Supernatant was removed, and the DNA pellet was dried at room temperature (RT) for 2–5 min. Rehydration of DNA in 250 µL of 1 X Tris-EDTA (pH 8) was performed at RT on a rotator for 20 RPM. overnight. Samples were stored at 4 °C for 2 days before use.

Cas9-targeted barcoding, library preparation, and long read sequencing

To perform targeted sequencing of *FMR1,* we designed and synthesized CRISPR–Cas9 crRNAs targeting the genomic regions adjacent to the *FMR1* CGG STR with the ChopChop online tool (**Table S1**). Preparation of the Cas9 nucleoprotein complex (Cas9 RNPs) was performed as follows: lyophilized Cas9 crRNA and tracrRNA (IDT) were suspended at 100 µM in TE (pH 7.5). The 4 crRNA probes (**Table S1**) were pooled for the cleavage reaction by combining equal volumes of each crRNA probe (0.25 µL each) and 1 µL tracrRNA (100 µM stock) in 8 µL of ultrapure molecular biology grade water. The pooled crRNAs and tracrRNA were annealed with a thermal cycler at 95° C for 5 min, allowed to cool to room temperature, and spun down to collect any liquid in the bottom of the tube.

To form Cas9 RNPs (for 10 reactions), components were assembled in a 1.5 mL Eppendorf DNA LoBind tube in the following order: annealed 10 µL crRNA•tracrRNA pool (10 µM), 10 µL 10 X NEB CutSmart buffer, 79.2 µL Nuclease-free water, and 0.8 µL HiFi Cas9 (62 µM, IDT). The tube was mixed thoroughly by flicking. RNPs were formed by incubating the tube at room temperature for 30 mins and then incubated on ice. Dephosphorylated genomic DNA was prepared by assembling the components in a 1.5 mL Eppendorf DNA LoBind tube in the following order: 5 µg of high molecular weight DNA in 24 µL, 3 µL NEB CutSmart Buffer (10 X), and 3 µL of QuickCIP enzyme (NEBM0525S). The sample was then incubated in a thermocycler at 37 ºC for 20 min, 80 ºC for 2 min, and held at 20º C. The reaction was then mixed gently and spun down.

Next, 10 µL of assembled RNPs from the previous step were incubated with 5 µg of dephosphorylated high molecular weight DNA, 1 µL of 10 mM dATP, and 1 µL of Taq polymerase (NEB) for 60 min at 37 °C on a thermocycler, followed by 5 min at 72 °C. 1 µL Proteinase K (Sigma, 20 mg/mL stock concentration) was added to each reaction, and samples were incubated at 43 °C for 30 min to remove proteins prior to size selection. The reaction was then purified to remove high concentration salt as follows: Cas9-cut genomic DNA (total volume is 42 uL) was precipitated using 16 uL of 5 M ammonium acetate together with 126 uL of ice-cold ethanol (absolute) and gently inverted ten times. Precipitated DNA was spun down at 16,000 g for 5 min. DNA was washed with 500 uL of 70% ethanol and centrifuged at 16,000 g for 5 min, and this step was repeated two times. The supernatant was removed, and the DNA pellet was dried at RT for 2-5 min. Rehydration of DNA was performed at 50 °C for 1 hr using 200 µL of 10 mM Tris-HCl (pH 8). DNA was further homogenized on a rotator at 37 °C and 20 RPM overnight. Size selection was then performed with the Bluepippin BLF7510 (Sage Science) using the "0.75DF 3-10kb Marker S1" cassette definition with size range at 5-12 kb.

To perform barcode ligation, 3 μL unique barcode (ONT EXP-NBD104) was added to 50 μL of Blunt/TA Ligase Master Mix (NEB) for each sample. The reactions were incubated at RT for 10 min, spun down, and then put on a magnet. The beads were washed twice with 200 μL of freshly prepared 70 % ethanol without disturbing the pellet, and allowed to dry for 30-60 seconds. The remaining pellet was then resuspended in 16 μL nuclease-free water and incubated for 10 min at room temperature. The reaction was then placed on a magnet, and 16 μL of supernatant was removed into a clean 1.5 mL Eppendorf DNA LoBind tube. Samples were then quantified using a Qubit fluorometer and Qubit dsDNA HS assay kit (Thermo Fisher Scientific).

Adapters were then ligated by adding 20 μL NEBNext® Quick Ligation Buffer (NEB#E6056S), 10 μL NEBNext Quick T4 DNA ligase (NEB#E6056S), and 5 μL Adapter Mix (AMII) at room temperature in a separate 1.5 mL Eppendorf DNA LoBind Tube. The ligation reaction was mixed thoroughly. 20 μL of the adapter ligation reaction was mixed with the pooled native barcode-ligated samples. Immediately after mixing, the remaining 15 μL of the adapter ligation mix was added to the native barcode-ligated sample to yield a 100 μL ligation mix. The reaction was incubated for 10 min at room temperature. Then 1 volume (100 μL) of TE (pH 8.0) was added to the ligation mix, followed by 0.4 X volume (80 μL) of AMPure XP Beads. The sample was then incubated for 10 min at room temperature, placed back on the magnet, and the supernatant was removed. The beads were then washed with 250 μL Long Fragment Buffer (LFB) twice and then air-dried for ~30 seconds. The library was eluted off the beads in 14 μL Elution Buffer. Finally, 13 uL of the library was then mixed with 37.5 μL sequencing buffer and 25.5 μL loading beads and loaded onto the MinION flowcell.

##### ***PCR Free Whole Genome Sequencing***

PCR Free whole genome sequencing libraries were aligned to hg19 using bwa-mem (v0.7.10-r789) and default parameters. Prior to mapping, reads were quality checked using FastQC (v0.11.9). After reads were aligned to hg19 using bwa-mem, the results were converted to .bam files and sorted using Samtools (v1.11). Using deeptools (v3.30) and Samtools flagstat, .bam files were quality checked, and mapping statistics were generated before proceeding to downstream analyses.

***Pre-mutation CGG STR transgene overexpression***

CGGx99 vector construction

A vector containing 99 CGG repeats within the *FMR1* 5'UTR was purchased from Addgene (63091). The CMV promoter in this vector was replaced by EF1a promoter. Briefly, the CMV promoter in the vector was removed by EcoRI and SalI digestion and replaced with a short fragment that contained two restriction cloning sites SpeI and BsiWI. The short fragment was generated from annealing two short oligos (5'-AATTCACTAGTGAATTCAGATCTGGTACCGTACG-3'; 5'-TCGACGTACGGTACCAGATCTGAATTCACTAGTG-3'). The EF1a promoter was isolated from another vector (Addgene, 104372) with NheI and BsiWI digestion and inserted to the CGG vector within SpeI and BsiWI restriction sites. The resulting construct generated the new expression vector EF1a-(CGG)x99-GFP.

CGGx99 vector transfection

All human iPSC lines were cultured in a 10 cm dish. CGG vector transfection was carried out with Lipofectamine stem reagent (ThermoFisher, STEM00008) by following the vendor's instruction. 24 hrs after transfection, cells were trypsinized and brought to the Children's Hospital of

Philadelphia flow core for sorting for both GFP negative and GFP positive cells. Sorted cells were continued in culture for another 24 hrs. The cells were then pelleted and used for qRT-PCR and CUT&RUN experiments.

***CUT&RUN***

CUT&RUN was completed as previously described (*12*). In brief, 300k-600k iPSC were washed in phosphate-buffered saline (PBS) and harvested 24 hrs after sorting (see: CGGx99 vector transfection). Harvested cells were then washed in wash buffer (20 μM Hepes KOH pH 7.5, 150 μM NaCl, 0.5 μM Spermadine, 1 Roche Complete Protease Inhibitor EDTA-free mini tablet per 10 mL) and bound to Concanavalin A beads (BioMagPlus) that had been activated and washed with binding buffer (20 μM Hepes KOH pH 8.0, 10 μM KCl, 1 μM $CaCl_2$, 1 μM $MnCl_2$). The cells were then incubated with the Concanavalin A magnetic beads, primary antibody (either IgG (Sigma I8140) or H3K9me3 (Abcam ab 8898)), and antibody buffer (digi-wash buffer – 0.1 % digitonin in wash buffer – with 2 μM EDTA) overnight at 4 °C. Cells were washed with digi-wash buffer and then incubated in a solution containing protein A-MNase and digi-wash buffer for 1 hr at 4 °C. After incubation, the samples were washed in digi-wash buffer, and 100 μL digi-wash buffer was added to the samples, which were then placed on an ice block sitting in an ice bath to chill for 5 min. After chilling, 2 μL of 100 μM $CaCl_2$ was added to activate protein A-MNase chromatin digestion. After 30 min, 100 μL of 2 X stop buffer (340 μM NaCl, 20 μM EDTA, 4 μM EGTA, 0.05 % Digitonin, 50 μg/mL RNase A, 50 μg/mL Glycogen) was added to halt the reaction, which was then incubated at 37 °C for 30 min to release chromatin fragments. Supernatant was collected, and DNA was extracted using phenol:chloroform and ethanol precipitation. The resulting DNA was quantified on a Qubit Fluorometer using Qubit dsDNA HS assay kit (Thermo

Fisher Scientific), and NEBNext Ultra II Library Prep Kit (NEB, E7645S) was performed using CUT&RUN specific PCR parameters as suggested by the EpiCypher CUTANA™ CUT&RUN protocol to selectively amplify fragments of interest. Fragments were characterized using Qubit Fluorometer with the dsDNA HS assay kit and Agilent Bioanalyzer 2100 (Agilent, 5067–4626) with the Agilent Bioanalyzer High Sensitivity DNA Analysis Kits (Agilent, 5067-4626). Libraries were pooled and paired-end sequencing was performed using the Illumina Nextseq 500 with the Ilumina Nextseq 500/550 High Output Kit v2 (75 cycles).

#### Data analysis

##### *Nanopore data processing for CGG tract analysis*

MinION sequencing reads were first processed using the base calling tool guppy_basecaller (Version 4.0.15). Then the base called reads were sorted by guppy_barcoder (Version 4.0.15) into each barcoded sample respectively. Reads were then corrected with canu (version 2.1.1) using default parameters. Reads that cover the *FMR1* loci were identified by extracting reads that contained a sequence upstream of the gene "GGAGGGAACAGCGTTGATCACGTG." All such reads covering the *FMR1* locus where the sequencing was done on the reverse orientation were extracted and used for further analysis. For each read, the following four characteristics was determined: (1) The total number of CGGs present. This was performed by counting the number of "CGG" instances between the start of the CGG tract and the end of the 5'UTR. (2) The longest continuous CGG track. This was determined by splitting the CGG tracts every time an "AGG" occurred and counting the number of CGGs in the resulting tracts. (3) The number of AGG interrupters within the CGG STR. This was determined by counting the number of "AGG" instances within the CGG tract. (e) the total number of continuous CGG tracks. This was determined by splitting the CGG tracts every time an "AGG" occurred and counting the number of resulting tracts. These four characteristics were calculated for every Nanopore long read that mapped to the *FMR1* loci and plotted.

##### *Nanopore data processing for DNA methylation analysis*

We basecalled long-reads using guppy (version 3.0.3+7e7b7d0) with configuration file "dna_r9.4.1_450bps_hac.cfg". Fastq files were preprocessed and resquiggled by tombo (version: 1.5.1) as described (https://github.com/bioinfomaticsCSU/deepsignal). Because each genotype has different CGG tract lengths, alignment of reads to hg19 yields inaccurate estimates. Hence, we edited the *FMR1* gene in-silico to include different CGG lengths, including 20 (WT_15), 180 (Prem_133 & FXS_326_CUT_180), 450 (FXS_326, FXS_306 & FXS_378), and 4 (FXS_378_CUT_4) CGG triplets. We next generated CpG methylation calls for each long read using DeepSignal (version 0.1.8) with mode "call_mods" and model "model.CpG.R9.4_1D.human_hx1.bn17.sn360.v0.1.7+". DeepSignal's script was used to call modification frequency "call_modification_frequency.py" at the reference genomic location. Finally, we plotted methylation frequency CpG positions with at least 3 reads mapped.

##### *ChIP-seq mapping*

ChIP-seq data was processed as previously described (*1, 4-9*). In brief, 75 bp single end reads were mapped to the hg19 reference genome using Bowtie with parameters: --tryhard -m 2. Optical and PCR duplicates were removed using samtools. Reads were downsampled to achieve equal read numbers across samples being compared (**Table S4**). CTCF peaks were called using MACS2 with a cutoff of p <$1x10^{-8}$ (**Table S7**). H3K9me3 domains were called using RSEG (see: H3K9me3 domain calling).

##### *5C analysis*

5C data was processed as previously described (*1, 4, 6, 7, 13-15*). In brief, 37 bp paired-end reads were mapped to a pseudo-genome consisting of all possible 5C primer ligation junctions with Bowtie using the following parameters: --tryhard and -m 2 and --trim5. All 5C primer-primer counts were represented as 2-dimensional matrices of interaction frequencies between each pairwise combination of primers. Outlier entries in the matrices, those which were 8-fold greater than the local media of the 5 surrounding entries, were filtered out. The interaction frequency matrices corresponding to samples to be compared were then quantile normalized together. The primer-primer interaction frequencies were then converted to fragment interaction frequencies as described previously (*6*). The fragment interaction frequencies were then binned into 4 kb resolution pixels, and a 6 kb smoothing window was applied to attenuate spatial noise. The binned and smoothed matrices were balanced using the ICED algorithm.

##### *Hi-C Data Processing*

Paired-end reads were aligned independently to the hg19 human genome using bowtie2 (global parameters: --verysensitive –L 30 –score-min L,-0.6,-0.2 –end-to-end --reorder; local parameters: --very-sensitive –L 20 –score-min L,-0.6,-0.2 –end-to-end --reorder) through the HiC-Pro software version 2.7.7[8]. Unmapped reads, non-uniquely mapped reads, and PCR duplicates were filtered, and uniquely aligned reads were paired. Raw cis contact matrices for all samples were assembled into 10kb, 20kb, 40kb, and 100kb non-overlapping bins and balanced using the Knight-Ruiz algorithm. The balanced cis matrices were then normalized across samples being directly compared using median-of-ratios size factors conditioned on genomic distance as we have previously described (*16*). Trans m x n contact matrices were assembled using Juicer by binning hg19 aligned, in situ Hi-C paired-end reads into uniform 1Mb-sized bins and balancing using the Knight Ruiz algorithm with default parameters. Trans matrices were quantile normalized across samples to facilitate direct comparison.

***CUT&RUN Data Processing***

Sequencing data was analyzed using Bowtie2 (version 2.2.5) with parameters "--local --very-sensitive-local --no-unal --no-mixed --no-discordant --phred33 -I 10 -X 700". Duplicates and unmapped reads were removed using Samtools (version 1.11) markdup command. After removing duplicates and unmapped reads, files were converted to bam files using Samtools, and then the resulting bam files were converted to bigwig format using BamCoverage from Deeptools (version 3.3.0). The "--normalizeUsing RPKM –extendReads" parameters for BamCoverage were used.

***Gene expression analysis – RNA-seq***

RNA-seq reads were mapped to the hg19 ensembl reference transcriptome for both cDNA and ncRNA using kallisto quant with 100 bootstraps of transcript quantification (*17*). Reads were mapped to the ensembl cDNA and ncRNA transcriptomes as described in the kallisto documentation. The resulting quantifications were converted into DESEQ2 format, and transcript level counts were mapped to gene level counts in R using the library("tximportData") according to DESEQ2 documentation recommendations (*18*). Genes with total counts less than 60 across all samples were dropped from analysis. Differentially called transcripts across the 5 cell lines studied were determined in a pairwise manner using DESEQ2 LRT with adjusted $p < 0.005$ (**Table S8**).

##### *H3K9me3 domain calling*

H3K9me3 domains were computationally identified using the RSEG program (version 0.4.9) (*19*). RSEG was run with parameters -s 400000 and with -d, deadzone flag, using RSEG provided deadzones for hg19. From the full list of domains calls, domains within 500 kb of centromeres were removed, and then domains located within 10 kb of each other using BedTools v2.29.2 were merged. In order to focus our analysis on large H3K9me3 domains, we filtered the full list of domains to for those greater than 200 kb in size. (**Table S6**). When RSEG domain calls were interrupted by unmappable regions with 0 mapped reads from H3K9me3 ChIP-seq data, the RSEG domains flanking the unmappable region were merged. We defined "Genotype Invariant H3K9me3 domains" as those present in 4/5 of WT, Premutation, short mutation-length FXS, and long mutation-length FXS lines, where RSEG domain calls had to have boundaries within 300kb of each other to be considered the same domain (**Table S5**). We defined "FXS-specific H3K9me3 domains" (total n=12) as those present in both long mutation-length FXS lines and not present in WT or invariant domains (**Table S5**).

##### *Insulation score calculation*

A 500 kb square window (50 x 50 bins on 10 kb binned data) with one bin offset from the diagonal was tiled across the genome on Knight-Ruiz balanced cis Hi-C maps. Counts in the 50 x 50 bin window were summed, normalized by the chromosome-wide mean, log transformed, and recorded as the Insulation Score (IS).

##### *Dimensionality index calculation*

To determine the directional bias of the bins corresponding to the genome locations of *FMR1*, the Directionality Index (DI) was used as described previously (*20*). Briefly, the directionality index is a weighted ratio between the number of Hi-C reads that map from a given 40 kb bin to the upstream region and the downstream region. 2 Mb upstream and downstream were used in the calculation.

##### *Compartment identification*

To determine A/B compartment status genome-wide, the eigenvector of the balanced, 100 kb binned cis Hi-C interaction matrix for each chromosome was calculated (*21, 22*). Briefly, the balanced matrix was first normalized by the expected distance dependence mean counts value, followed by removal of rows and columns that were composed of less than 2 % non-zero counts. The off-diagonal counts were then z-scored, after which a Pearson correlation matrix for the cis-interaction matrixes was calculated. The eigenvector was selected as the largest eigenvalue of the Pearson correlation matrix computed from the Hi-C matrix. The coordinates corresponding to transitions between positive and negative eigenvector values demarcate boundaries of compartments. Using the established pattern of gene density in A/B compartments, we assigned positive eigenvector values to the gene-dense A compartment, and negative values to the gene-poor B compartment.

##### *Binning ChIP-seq & A/B compartment signal*

Binned H3K9me3 signal shown in Figure 1F was generated by taking the H3K9me3 ChIP-seq signal across the loci of interest, splitting the loci into 40 evenly sized bins, and plotting one point for the average ChIP-seg signal of each bin. Similarly, compartment score in Figure 1G was calculated by splitting the locus of interest into 40 evenly sized bins, and plotting one point for the average compartment score of each bin. For Figure 2A, H3K9me3 signal was plotted in heatmap form for "genotype-invariant H3K9me3 domains", "FXS-specific H3K9me3 domains consistently gained in both long mutation-length FXS lines", and "FXS-specific H3K9me3 domains gained in only one FXS line" by binning ChIP-seq signal into 100 equally sized bins. To scale all the domains, which are different sizes, to be represented as the same width in the heatmaps, the average H3K9me3 ChIP-seq signal in each bin was calculated and plotted. The flanking 100 kb regions around each domain were also binned into 100 equally sized bins, and the average H3K9me3 ChIP-seq signal in each bin was calculated and plotted.

***Identification of genes in H3K9me3 domains***

Genes were defined as co-localized to H3K9me3 domains if the TSS of the gene was contained within the domain. The intersections were performed using BedTools.

***Quantifying long-range interaction frequency among key genes from Hi-C***

To determine the contact frequency between *FMR1* and *SLITRK2,* normalized (see section Hi-C Data processing, above) Hi-C data binned at 20 kb resolution was used. The normalized counts in bins corresponding to interactions between the hg19 coordinates of *FMR1* and *SLITRK2* in the cis chr X interaction matrix were summed. To determine the contact frequency between *FMR1* and *SLITRK4,* normalized (see section Hi-C Data processing, above) Hi-C data binned at 40 kb resolution was used. The normalized counts in bins corresponding to interactions between the hg19 coordinates of *FMR1* and *SLITRK4* in the cis chr X interaction matrix were summed.

##### *CTCF motifs*

The location of CTCF motifs in hg19 was obtained from the JASPER database using the following parameters: hg19 reference genome, JASPER 2018 consensus, motif: CTCF, allow overlapping motifs, pvalue = 0.001, search both strands.

##### *Ideograms and domain location*

Ideograms were retrieved from the UCSC genome browser by using the UCSC Table Browser for hg19 and selecting Group="All Tables" and Table="cytoBand". The location of the red boxes corresponding to gained H3K9me3 domains in FXS were determined by using the UCSC genome browser to locate the coordinates on the ideogram.

##### *Gene ontology analysis*

Gene ontology enrichment was performed using WebGestalt (http://www.webgestalt.org) with the following settings: Organism of interest = homo sapiens; Method of interest = overrepresentation enrichment, Functional database = geneontology, biological_process_noRedun. Gene name identifiers were uploaded for each set of classified genes. The genome_protein-coding set was used as the reference set. The enrichment ratios and −log10(p-values) for all gene ontology terms with an p of $< 0.01$ and enrichment ratio $> 4$ were plotted.

The input gene lists for gene ontology analysis were determined as follows: In Figure 2i, all genes which had their TSS reside in “FXS-specific H3K9me3 domains consistently gained in both long mutation-length FXS lines" (see above: "H3K9me3 domain calling"), had expression greater than 0 in at least one of the cell lines where RNA-seq was performed, and were protein coding (microRNAs and long non coding RNAs were excluded) were input into WebGESTALT. Only protein coding genes were included using the genome_protein-coding set as the reference set. In Figure 2j, genes were selected in a similar manner, but using FXS-specific H3K9me3 domains variably gained in only one long mutation-length FXS line" (see above: "H3K9me3 domain calling").

##### *GTEX Gene Expression Data*

Gene expression across human tissues was obtained from the GTEX consortium. The data used for the analyses described in this manuscript were obtained from https://www.gtexportal.org/home/datasets on the GTEx Portal on 04/2020. To generate the heatmap in Figure 2h, the expression of all genes in n=12 consistently gained H3k9me3 domains in FXS was first retrieved. Then, genes which had 0 expression across all tissues were removed, resulting in a final list of n=67 genes. Then, gene expression data was z-scored across tissues (such that strong expression of one gene in one tissue type does not wash out signal in all other tissues). Finally, genes were clustered on the gene expression data using K-means clusters into 4 groups. Clusters were labelled based on the tissue types dominating each cluster.

##### *Location of CGG short tandem repeat tracts in hg19*

Location of CGG repeats in hg19 were identified by string search from the hg19 reference genome. Any strings of more than two CGGs in a row were included in the analysis.

##### *Enrichment tests for annotations in FXS specific domains*

To determine whether unstable STRs are enriched in FXS specific H3K9me3 domains (test set) compared to the genome-wide distribution, the total number of unstable STR tracks (see below: STR genotyping and instability analysis) located in genes within the test set was summed. Then, the same sum of total number of unstable STR tracks was determined for 1000 random draws of n=12 size matched genomic intervals (null set). An empirical p-value was calculated to determine which set was enriched for unstable STRS in genes compared to the null set. To determine whether STRs are enriched in FXS specific H3K9me3 domains (test set) compared to genotype invariant domains, this process was repeated using an alternative null set consisting of 1000 random draws of n=12 size matched intervals centered over genotype invariant H3K9me3 domains.

To determine whether fragile sites are enriched in FXS specific H3K9me3 domains (test set) compared to the genome-wide distribution or to genotype invariant domains (null sets), the same method as for unstable STRs was performed but counting the total number of fragile sites within either the test or null set instead. Fragile sites were obtained from the HumCFS database.

##### *Identification of FXS specific H3K9me3 domain as either reprogrammed vs resistant to CGG deletion*

FXS specific H3K9me3 domains were categorized as either reprogrammed or resistant to CGG deletion based on presence of the domain in the FXS_iPSC_326 iPSC line compared to in an isogenic line where the CGG tract was cut from >450 to 180 (FXS_326_CUT_180, see above for methods). Resistant H3K9me3 domains persisted despite CGG deletion and are present in both iPSC and FXS_326_CUT_180 as determined by RSEG domain calling (see above); reprogrammed domains were lost in FXS_326_CUT_180.

***STR genotyping and instability analysis***

We aligned PCR-free whole genome sequencing data of comparable sequencing depth (~ 450 million reads per sample, **Table S10**) to hg19 using bwa-mem (version 0.7.10) with default parameters (see full methods above).

We ran GangSTR (version 2.5.0) with the STR input file "hg19_ver13_1.bed" from GangSTR github page and a custom STR file, which we generated that included locations of CGG/CCG (>=2 consecutive units) genome-wide in hg19 (see methods above). Default parameters with one additional parameter declaring sex as males (--samp-sex M) were used. The resulting data consisted of the length of each STR loci for WT_15, Prem_133, FXS_306, FXS_326, and FXS_378 (**Table S9**). If more than one alternate allele was present, we took the length of the longest allele. Scatter plots (i.e., dot plots) of length were made using the default function 'plot' in R (version 4.0.3).

We formulated a statistical test to ascertain if normal-length STRs which grow unstable in FXS would be enriched in our FXS-specific H3K9me3 domains as compared to the expected instability at across the human genome. Our null hypothesis was that normal-length STRs which show instability (expansions or contractions) in FXS would be randomly distributed across the genome. By contrast, our alternative hypothesis was that normal-length STRs which show instability (expansions or contractions) in FXS would be significantly enriched at FXS-specific H3K9me3 domains compared to genotype-invariant H3K9me3 domains or genome-wide, size-matched, random locations. We first filtered the starting STR list (N = 15, 7323 STR sequence types; N = 966,459 total STR tracts) for only those STR classes in FXS specific H3K9me3 domains that expanded in at least one out of three disease lines (FXS_iPSC_306, FXS_iPSC_326, or

FXS_IPSC_378) by at least 3 repeat units compared to both wildtype (FXS_iPSC_15) and premutation (Prem_iPSC_133) lines, moving (N = 71 STR sequence types; N = 290 total STR tracts) unstable STR sequence types forward into the statistical test. We created empirical null distributions using two types of control regions, including random genomic locations size-matched to the "FXS-specific H3K9me3 domains" and genotype-invariant H3K9me3 domains size-matched to the "FXS-specific H3K9me3 domains".

We conducted a statistical test for every one of our N=71 STR classes. We formulated a test-statistic represented generally as:

$$T_{STR\_class} = \sum_{i=0}^{p} |STR\_tract\_length_{Disease} - STR\_tract\_length_{WT}|$$

where p is the total number of individual STR tracts within a particular STR sequence class. First, we computed the main test statistic as:

$$T_{STR\_class\_in\_FXS\_H3K9me3\_domains} = \sum_{i=0}^{n} |STR\_tract\_length_{Disease} - STR\_tract\_length_{WT}|$$

where n is the total number of individual STR tracts within a particular STR sequence class in our N=12 FXS-specific H3K9me3 domains. Next, we generated an empirical null distribution using the same test statistic:

$$T_{STR\_class\_in\_random\_size\_matched\_domains} = \sum_{i=0}^{m} |STR\_tract\_length_{Disease} - STR\_tract\_length_{WT}|$$

where m is the total number of individual STR tracts within a particular STR sequence class in each of 1,000 random draws of N=12 size-matched, random domains genome-wide or N=12 size-matched, genotype-invariant H3K9me3 domains genome-wide. We computed a one-tailed P-value for the probability of observing T greater than or equal to $T_{STR\_class\_in\_FXS\_H3K9me3\_domains}$ given our null distribution of 1,000 $T_{STR\_class\_in\_random\_size\_matched\_domains}$ or $T_{STR\_class\_in\_size\_matched\_genotype\_invariant\_H3K9me3\_domains}$. After obtaining P-values for all N=71 STR classes, we multiple testing corrected with Benjamini-

Hochberg step-up. Only STR classes passing a False Discovery Rate threshold of 10% in both genome-wide random or genotype-invariant H3K9me3 domains were reported as unstable STRs in FXS in which the null hypothesis can be rejected (in favor of high enrichment of unstable STRs in FXS-specific H3K9me3 domains vs. a random genome-wide distribution).

Beyond specific STR sequences and tracts, we also formulated a statistical test to assess overall instability burden of total STR tracts in FXS-specific H3K9me3 domains compared to random genome locations. We formulated 2x2 contingency tables to assess if STRs which grow unstable in FXS (total of N = 290 independent STR tracts for N = 71 STR sequence classes) are more enriched in FXS-specific H3K9me3 domains compared to STRs which do not grow unstable (total of N = 7998 independent STR tracts). We computed an Odd's Ratio test statistic as OR= (a/b)/(c/d) using either N=12 size-matched, random domains genome-wide or N=12 size-matched, genotype-invariant H3K9me3 domains genome-wide. We used Fisher's Exact test to compute P-values, and conducted 100 permutations of these tests to evaluate the 95% confidence intervals around the Odd's Ratio and the average P-value across 100 tests.

| | N=12 FXS-specific H3K9me3 domains | N=12 Random, size-matched domains |
|---|---|---|
| STR tracts which grow unstable in 1/3 disease lines compared to 176/111 by at least 3 units | 290 | 213 |
| STR tracts which do not grow unstable in any of the 5 lines | 7998 | 9704 |

| | N=12 FXS-specific H3K9me3 domains | N=12 Genotype-invariant H3K9me3 domains |
|---|---|---|

| | | |
|---|---|---|
| STR tracts which grow unstable in 1/3 disease lines compared to 176/111 by at least 3 units | 290 | 154 |
| STR tracts which do not grow unstable in any of the 5 lines | 7998 | 6441 |

<u>**Supplementary Figures**</u>

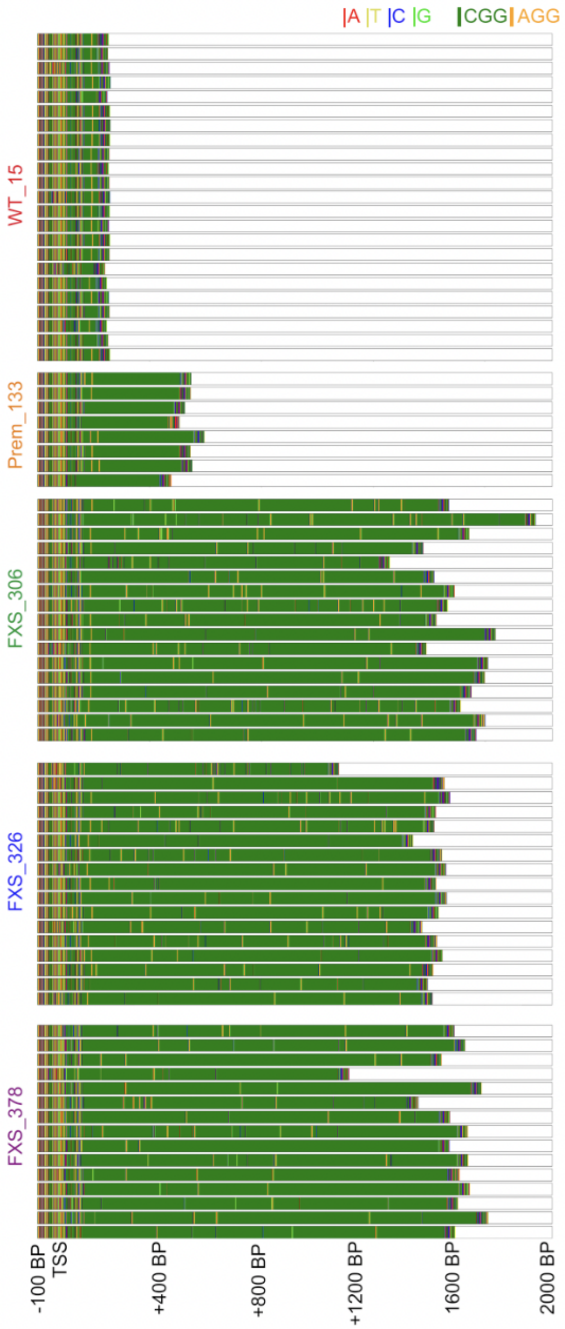


**Fig. S1. Nanopore long-read sequencing of the *FMR1* CGG short tandem repeat tract.**
Visual representation of Nanopore long reads that span the transcription start site and first 200 bp of human *FMR1*. For each of the 5 samples, the sequence of each read is shown with colors corresponding to base pairs as shown in the legend (top right). Only reads oriented on the reverse/antisense strand are shown.

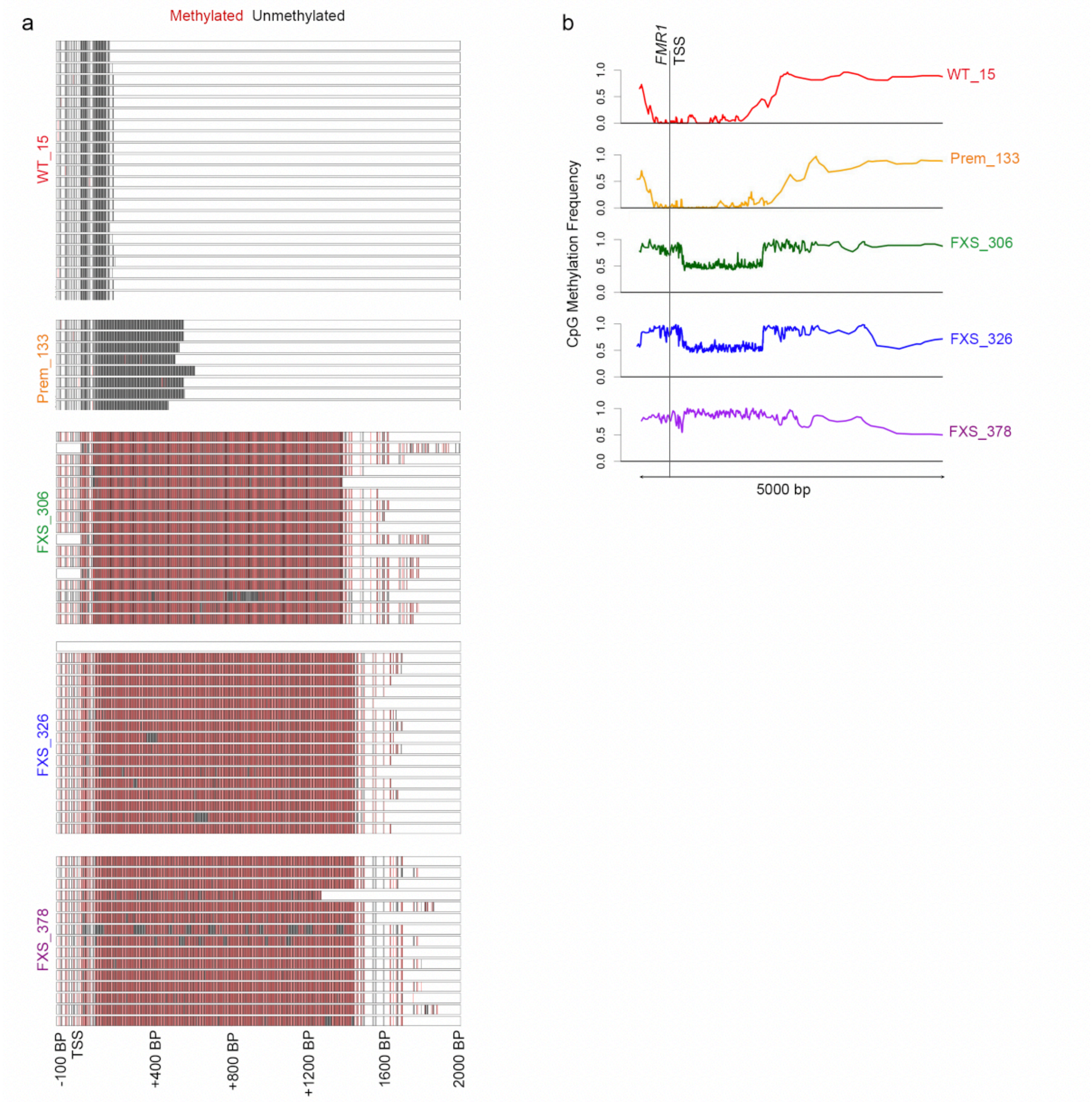


**Fig. S2. CpG methylation evaluated from Nanopore long-reads is increased over the *FMR1* transcription start site and CGG STR in fragile X syndrome.**

(a) For each long read obtained per cell line, DeepSignal was used to determine the methylation status of each CpG present in the sequence (see Methods). Black indicates an unmethylated CpG and red indicates a methylated CpG. Blanks indicate no CpG present. (b) For each cell line, DNA methylation for each read was averaged over all reads.

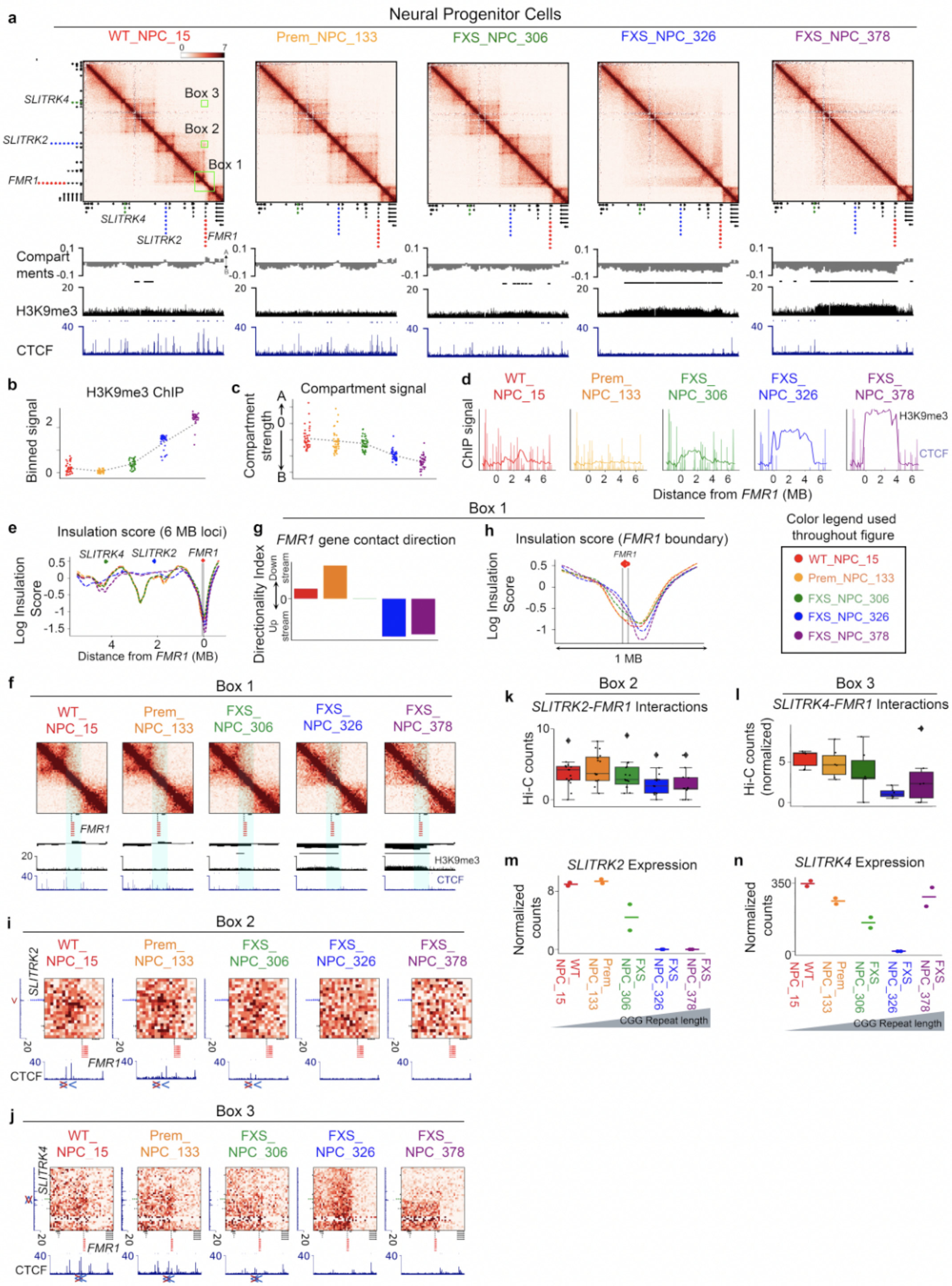


**Fig. S3. Genome folding disruption upon acquisition of a Mb-sized heterochromatin domain across the *FMR1* locus as the CGG STR tract expands from short mutation length to long mutation-length in fragile X syndrome.**
(a) Hi-C data across all five iPSC-NPC lines is shown as a heatmap of counts representing interaction frequency for an 8 Mb region around *FMR1*. A/B compartment score, H3K9me3 Chip-seq, and CTCF Chip-seq in iPSC-derived NPCs is displayed below the heatmaps for all

five conditions. Box 1, 2, and 3 are highlighted and referenced elsewhere in the figure. (b) H3K9me3 ChIP signal across the entire loci shown in panel (a) is binned into 40 bins and plotted for each cell line. Each dot represents one bin. (c) A/B compartment score across the loci shown in panel (a) is binned into 40 bins and plotted for each cell line. Each dot represents one bin. (d) CTCF and H3K9me3 ChIP-seq including *FMR1* and up to 6 Mb upstream are overlaid on each other for each cell line. (e) Insulation score up to 6Mb upstream from *FMR1* is shown for 5 cell lines. Grey vertical lines represent location of *FMR1* gene. (f) Zoom in on Box 1 from panel (a) on a 1 Mb region centered on *FMR1*. Blue highlights demonstrate location of gained contains/boundary disruption in disease. (g) Directionality index, a metric quantifying domain boundary strength, is plotted at *FMR1* in 5 cell lines. (h) Insulation score in a 1 Mb region containing *FMR1* is shown for 5 cell lines. Grey vertical lines represent location of *FMR1*. (i, j) Zoom-ins to Box2 and Box3 from panel (a) showing *FMR1-SLITRK2* or *FMR1-SLITRK4* gene-gene interactions. (k,l) Boxplot showing the interaction frequency measured with Hi-C between *FMR1* and either *SLITRK2* (k) or with SLITRK4 (l). Genomic intervals containing *FMR1, SLITRK2, SLITRK4* are binned into 20 kb bins and each dot represents interactions between one set of bins. A total of n=15 bins are shown for *SLITRK2* and n=6 bins are shown for *SLITRK4*. Boxes show the range from lower to upper quartiles, with median line, and whispers extend to minimum and maximum data points with 1.5 times the interquartile range. (m, n) Expression of *SLITRK2* and *SLITRK4* across 5 cell lines. Data is shown for 2 replicates. Dots represent replicates in expression plot, and lines represent mean expression across replicates.

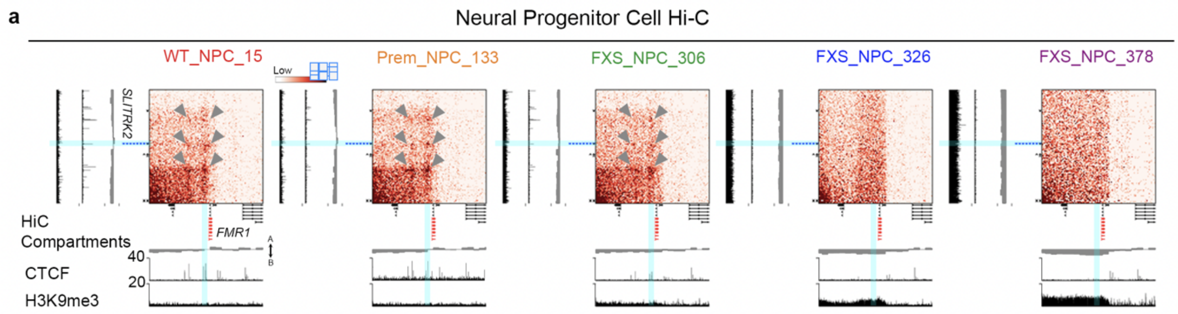


**Fig. S4. A series of loops connecting *FMR1* to *SLITRK2* are lost in fragile X syndrome**. Heatmaps of Hi-C cis interaction frequency in a 2 Mb window around *FMR1* (red, x-axis) interacting with a 2.6 Mb window around *SLITRK2* (blue, y-axis) across five iPSC lines differentiated to NPCs with normal-length, pre-mutation, short mutation-length, or long mutation-length CGG STR tract in *FMR1*. A/B compartment score computed from Hi-C data, CTCF ChIP-seq, and H3K9me3 ChIP-seq tracts are shown below heatmaps for each cell line. Grey arrows denote locations of loops which are lost in short and long mutation-length FXS lines. Turquoise bars highlight the location of CTCF sites that are lost in parallel with looping interactions

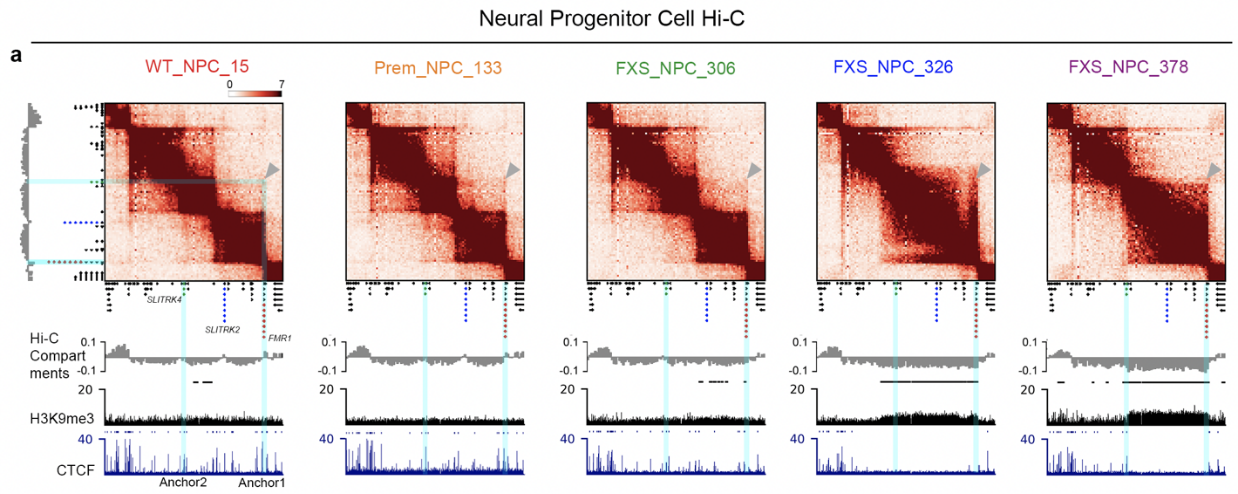


**Fig. S5. Two long mutation-length fragile X syndrome samples differ in the spread and density of H3K9me3 domain.**
Hi-C heatmaps of cis interaction frequency in a 10 Mb window around *FMR1* (red gene) *SLITRK2* (blue gene), and *SLITRK4* (green gene) across five iPSC lines differentiated to NPCs with normal-length, pre-mutation, short mutation-length, or long mutation-length CGG STR tract in *FMR1*. A/B compartment score computed from Hi-C data, CTCF ChIP-seq, and H3K9me3 ChIP-seq tracts are shown below heatmaps for each cell line. Grey arrows denote locations of long-range ~5 Mb loops between *FMR1* and *SLITRK4* in WT. Turquoise highlights over *FMR1* and *SLITRK4* are shown and intersect at the location of the grey arrow.

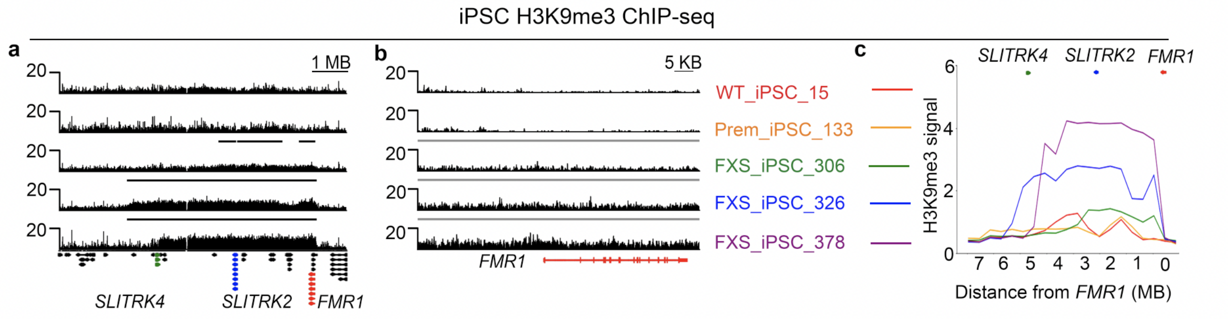


**Fig. S6. A megabase-scale heterochromatin domain is deposited across the *FMR1* locus in FXS iPSC**.

(a) H3K9me3 Chip-seq in one replicate of five iPSC lines in an 8 Mb region around *FMR1*. Genes are shown below ChIP-seq tracks. *FMR1*, *SLITRK2*, and *SLITRK4* are highlighted in red, blue, and green respectively. (b) Zoom-in on data from (a) is shown in an 75 kb window around the *FMR1* gene. (c) H3K9me3 ChIP-seq from (a) is shown with all 5 cell lines overlaid directly on top of each other.

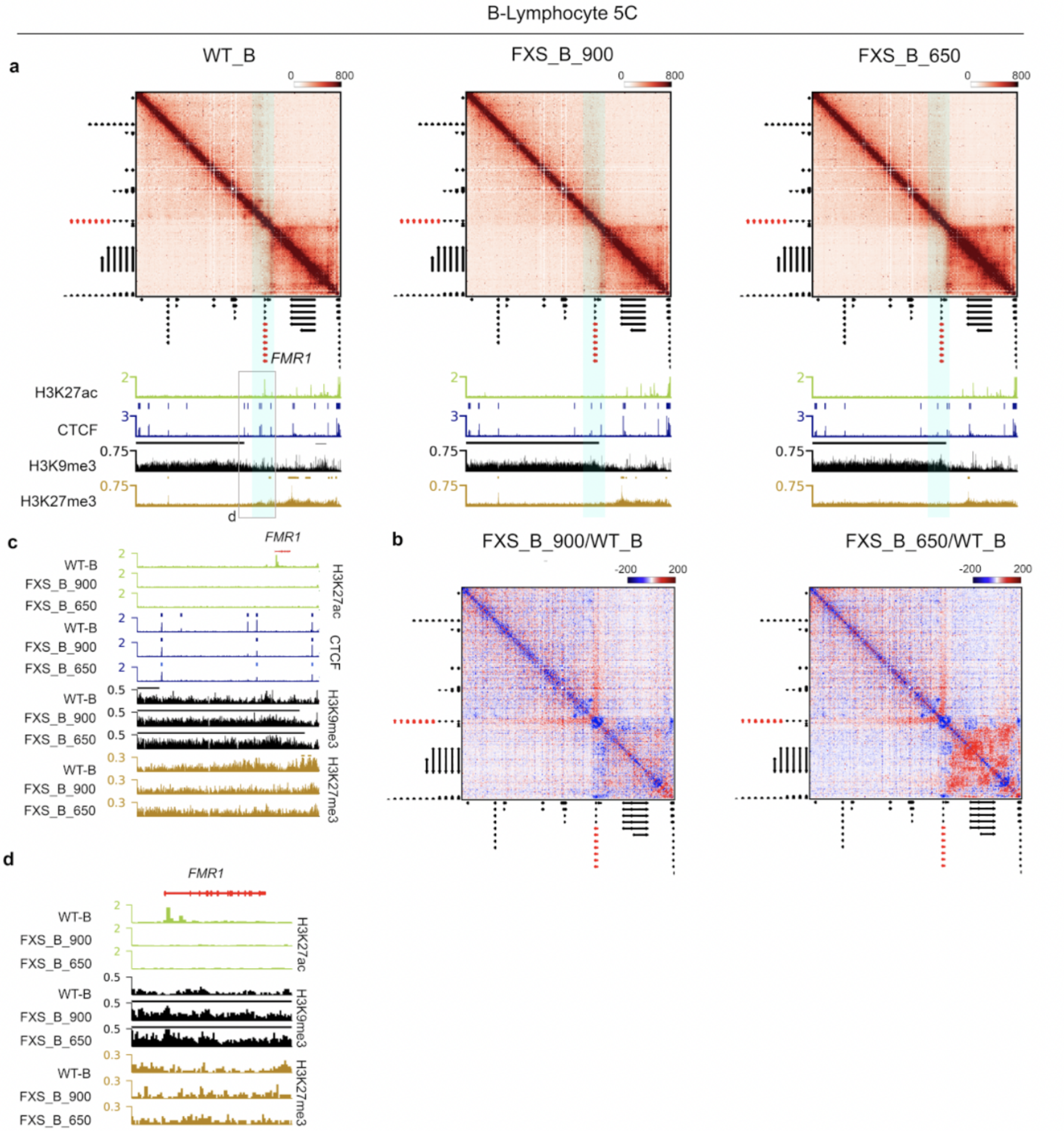


**Fig. S7. Local 3D genome alterations such as TAD boundary disruption and loop loss occur around the *FMR1* gene upon gene silencing and mutation-length CGG expansion.**
(a) 5C Heatmaps of ~5 Mb of the X chromosome surrounding the *FMR1* gene in WT B cells, B cells from FXS patients with 900 CGG repeats in the 5'UTR of *FMR1*, and B cells with 650 CGG repeats from a different patient. H3K27ac, CTCF, H3K9me3, and H3K27me3 ChIP seq tracks from these B-cells are aligned underneath heatmaps. All data is from 1 replicate. (b) 5C data in each FXS B-cell line is divided by 5C data in WT B-cells to show fold change maps aligned with epigenetic modification tracks in (a). (c) Zoom in on to 1.5 Mb around *FMR1* (location marked in grey rectangle in (a). (d) Zoom in on to 80 kb around *FMR1*.

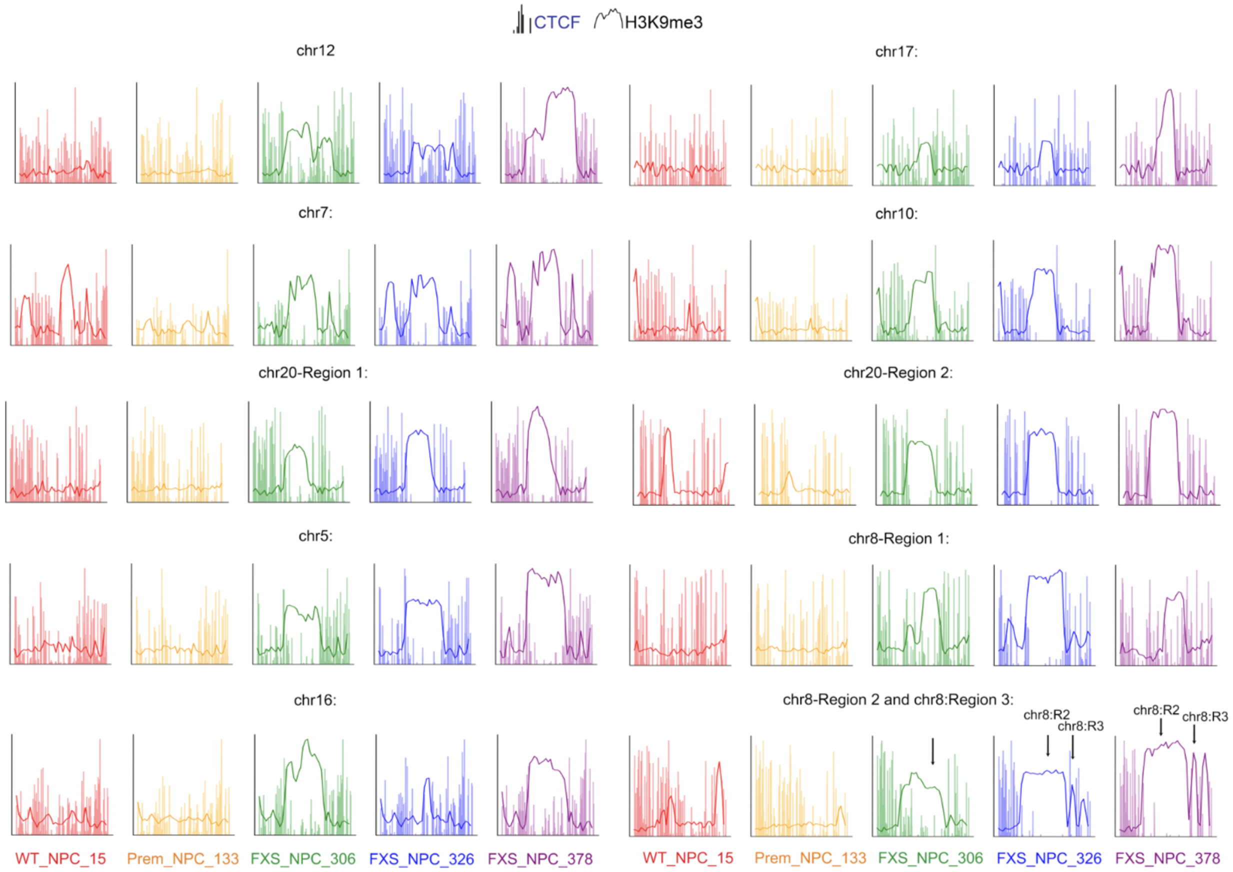


**Fig. S8. Distal FXS H3K9me3 domains in iPSC-derived NPCs.**
H3K9me3 and CTCF Chip-seq are plotted at n=11 distal FXS H3K9me3 domains in five NPC lines (normal-length (15 CGG), pre-mutation (133 CGG), short mutation-length (306 CGG), long mutation-length sample 1 (326 CGG), long mutation-length sample 2 (378 CGG)). Note: two of the 11 domains are separated by only 200kb, so both are shown on one plot in the chr8:134.6-142.7 interval (noted by arrows).

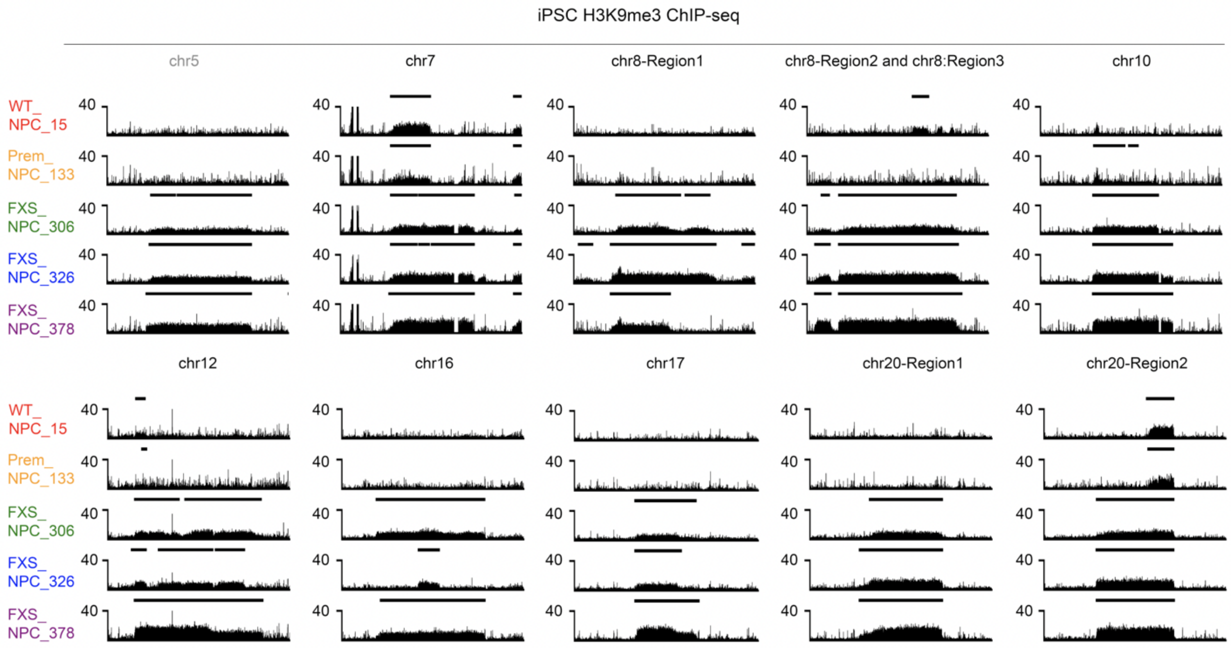


**Fig. S9. Distal FXS H3K9me3 domains in iPSC.**
H3K9me3 Chip-seq is shown around distal FXS H3K9me3 domains for five iPSC lines (normal-length (15 CGG), pre-mutation (133 CGG), short mutation-length (306 CGG), long mutation-length sample 1 (326 CGG), long mutation-length sample 2 (378 CGG)). Lines representing RSEG H3K9me3 domain calls are shown above H3K9me3 ChIP-seq track. Of n=12 total H3K9me3 domains gained, one is at *FMR1* (**Fig. S6**), and the remaining 11 are shown here. Note: two of the 11 domains are separated by only 200kb, so both are shown on one plot in the chr8:134.6-142.7 interval.

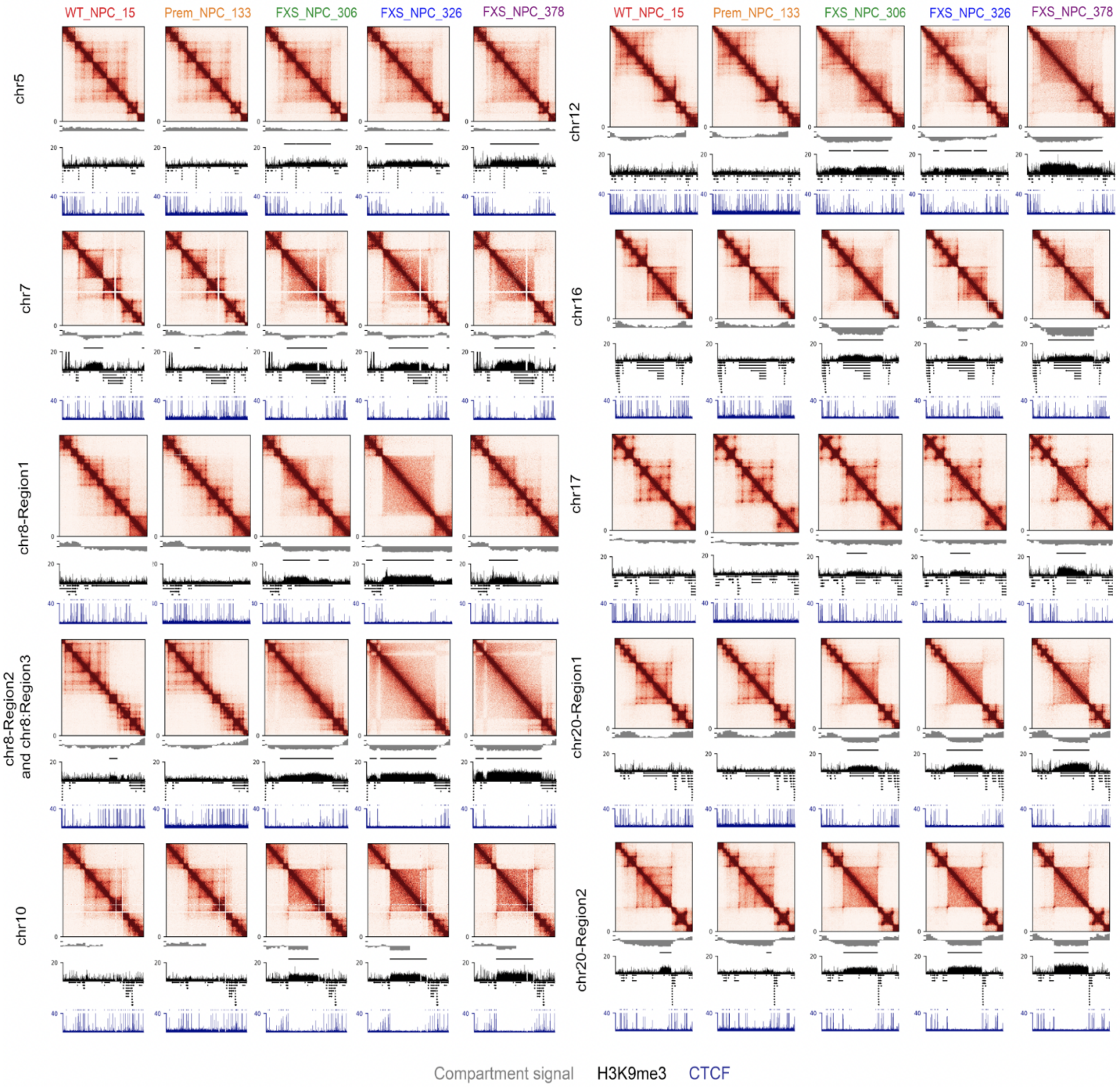


**Fig. S10. Genome folding is severely disrupted upon acquisition of distal H3K9me3 domains in FXS.**

Hi-C Interaction frequency heatmaps around distal FXS H3K9me3 domains. A/B compartment score, H3K9me3 ChIP-seq, and CTCF ChIP-seq from NPCs is displayed below the heatmaps. Lines representing RSEG H3K9me3 domain calls are shown above the H3K9me3 ChIP-seq track. Of n=12 total H3K9me3 domains, one is at *FMR1* (Figure 1e) and the remaining 11 are shown here.

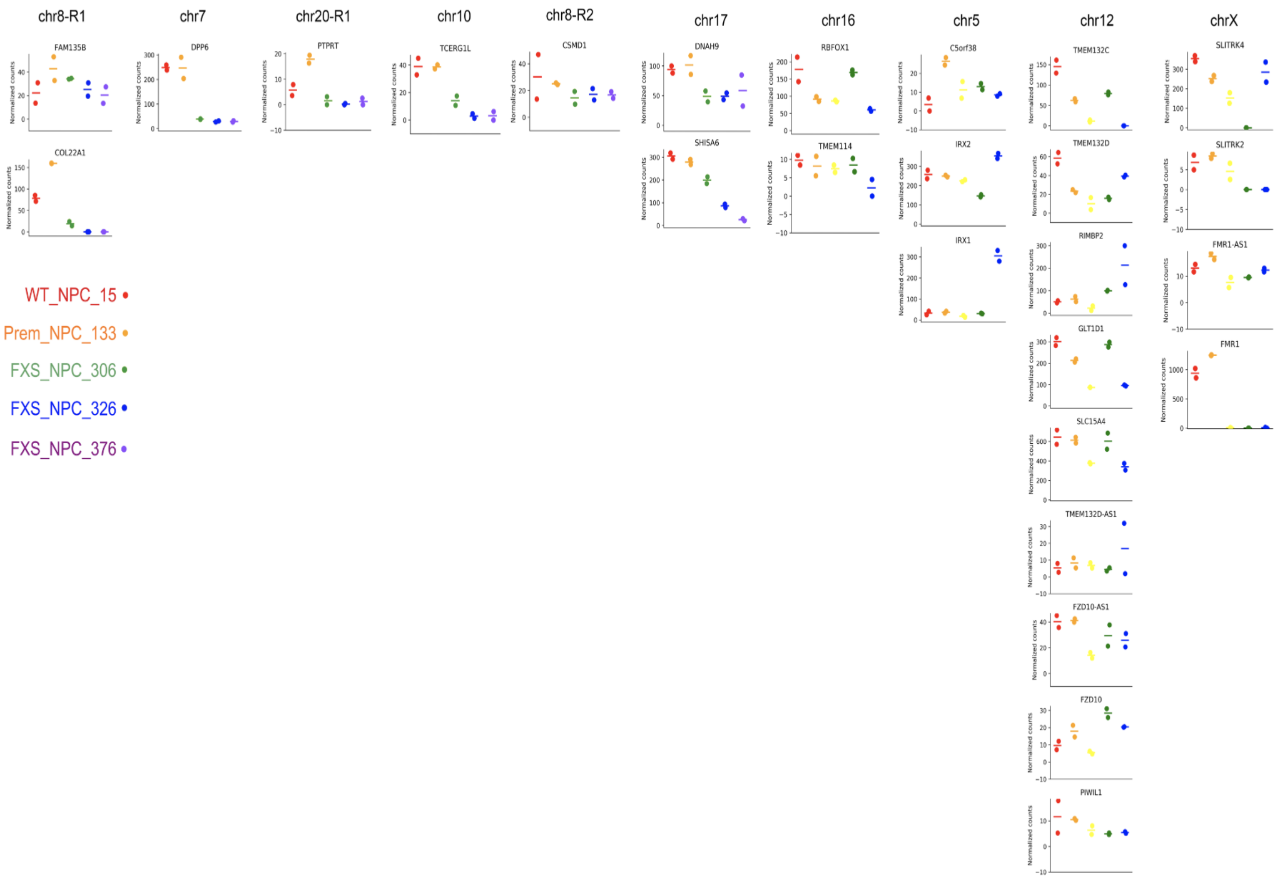


**Fig. S11. Expression of genes in n=12 FXS H3K9me3 domains.**
Of the n=12 domains, 10 contained protein coding genes expressed in NPCs. For each domain, expression for genes in that domain is shown for normal-length (WT_NPC_15), pre-mutation (Prem_NPC_133), short mutation-length (FXS_NPC_306), long mutation-length sample 1 (FXS_NPC_326), long mutation-length sample 2 (FXS_NPC_378). Two biological replicates are shown for each sample. Horizontal line represents the mean of two replicates for each of five lines.

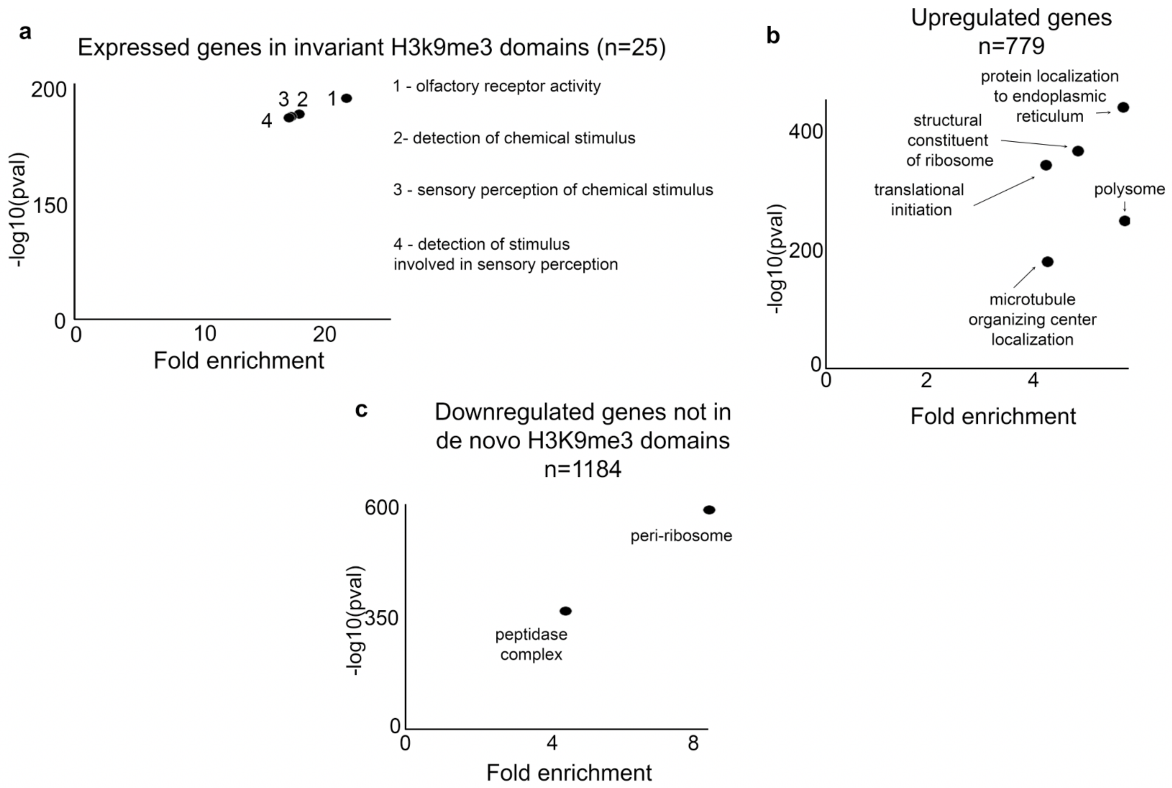


**Fig. S12. Gene Ontology for upregulated and downregulated genes in FXS.**
Gene ontology for (a) n=25 protein coding expressed genes in invariant H3K9me3 domains as defined in Figure 2a, (b) genes upregulated in both long diseases, or (c) genes downregulated in both long mutation-length FXS lines but not located within one of n=12 consistently gained FXS H3K9me3 domains. GO was performed using WebGESTALT with settings Over-Representation Analysis, geneontology, Biological Process, Cellular Component, Molecular function, with "genome-protein coding" as the reference. A P-value cutoff of $p < 0.01$ and enrichment > 4 was used.

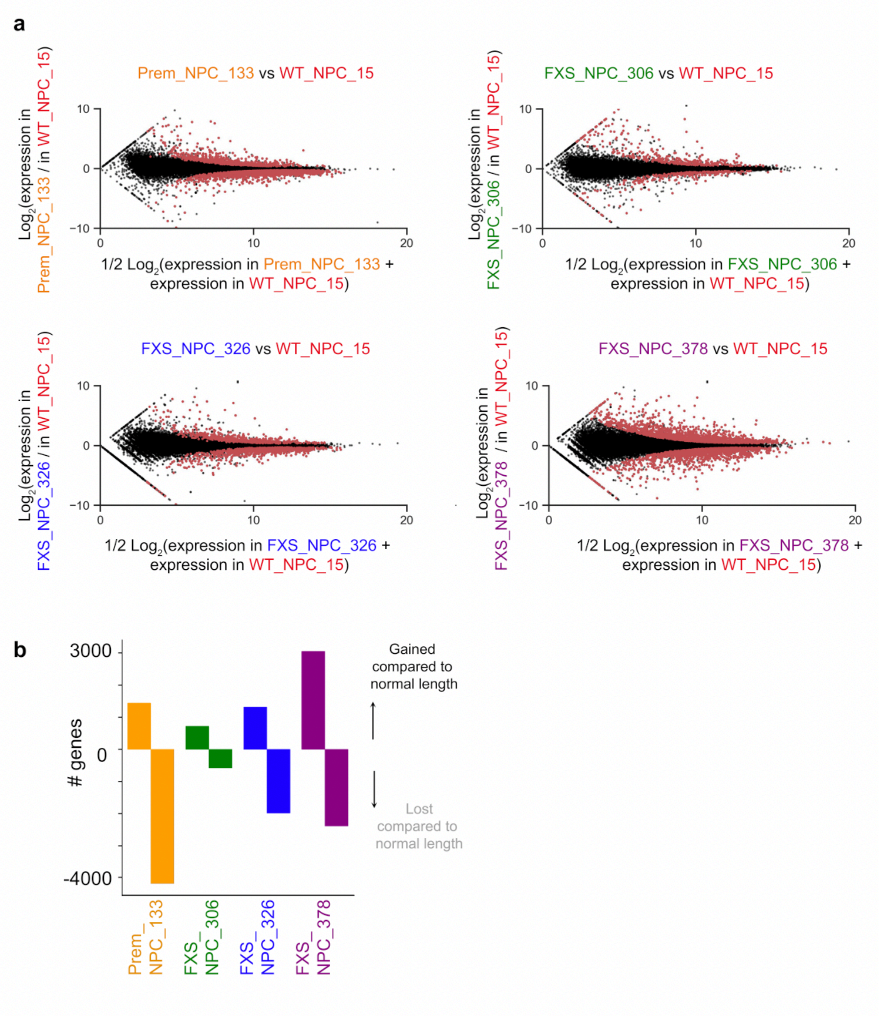


**Fig. S13. Differential gene expression across FXS cell lines.**
(a) M-A plot showing RNA-seq data in iPSC-derived NPCs (normal-length (WT_NPC_15), pre-mutation (Prem_NPC_133), short mutation-length (FXS_NPC_306), long mutation-length sample 1 (FXS_NPC_326), long mutation-length sample 2 (FXS_NPC_378)). Genes in red are called significant by DEseq2 using a likelihood ratio test at a threshold of $p < 0.005$. (b) Total number of up- and down-regulated genes for all lines (pre-mutation (Prem_NPC_133), short mutation-length (FXS_NPC_306), long mutation-length sample 1 (FXS_NPC_326), long mutation-length sample 2 (FXS_NPC_378)) compared to normal-length (WT_NPC_15).

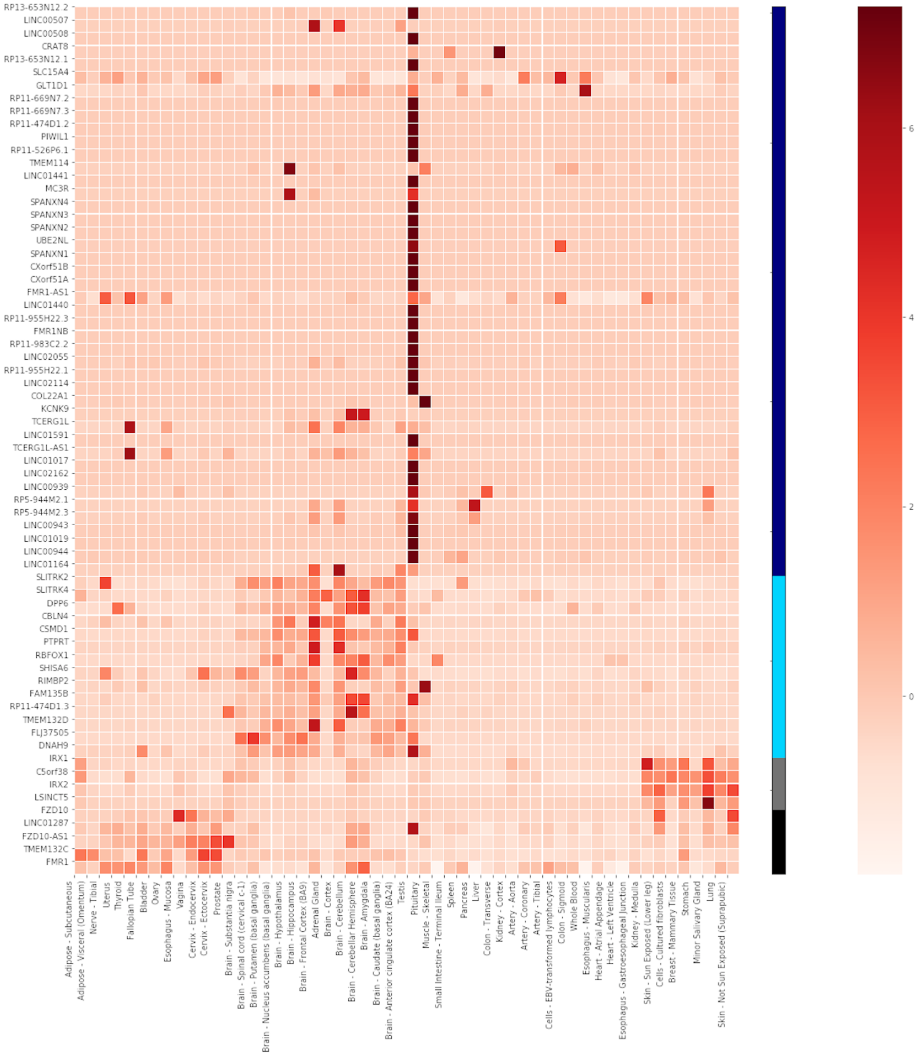


**Fig. S14. Tissue specific expression of genes in FXS H3K9me3 domains.**

Expression of all genes in H3k9me3 domains consistently gained across all 3 FXS lines in n=54 tissues in GTEX dataset. Genes were only shown if expression was not zero across all tissues, resulting in n=67 genes. Genes were clustered using K-means clusters into 4 groups, and clusters were labelled based on the tissue types dominating each cluster. This is the same as figure 2h, but with all labels for each axis shown legibly in a larger image footprint.

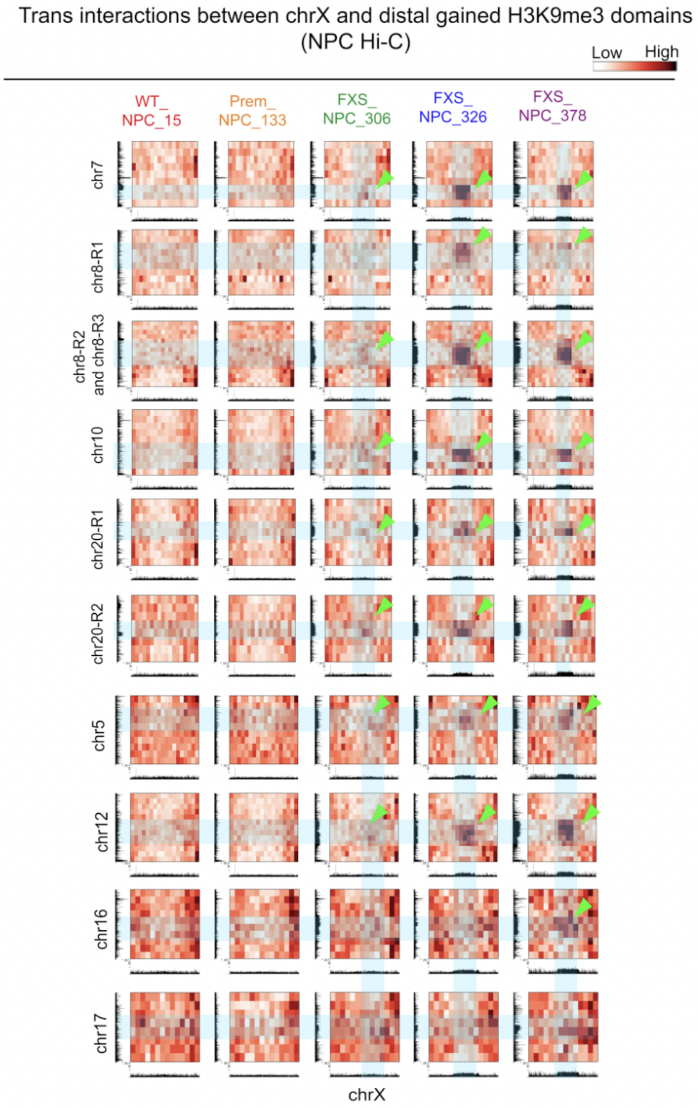


**Fig. S15. Inter-chromosomal interactions between *FMR1* and distal FXS H3K9me3 domains.**

Hi-C interactions between *FMR1* and each of the distal H3K9me3 domains gained are shown for normal-length (WT_NPC_15), pre-mutation (Prem_NPC_133), short mutation-length (FXS_NPC_306), long mutation-length sample 1 (FXS_NPC_326), and long mutation-length sample 2 (FXS_NPC_378) *FXS* NPCs. The window for each region includes the H3K9me3 domain gained and 5 Mb of flanking genome. H3K9me3 ChIP-seq *FMR1* is shown on the x axis (chr X) and for the distal region is shown on the Y axis. All data is from one replicate per cell line. Hi-C data is binned at 1 Mb resolution. The domains are identified by which chromosome they are on, and the *FMR1* domain is identified as “chrX.” Blue bars and green arrows highlight trans interactions.

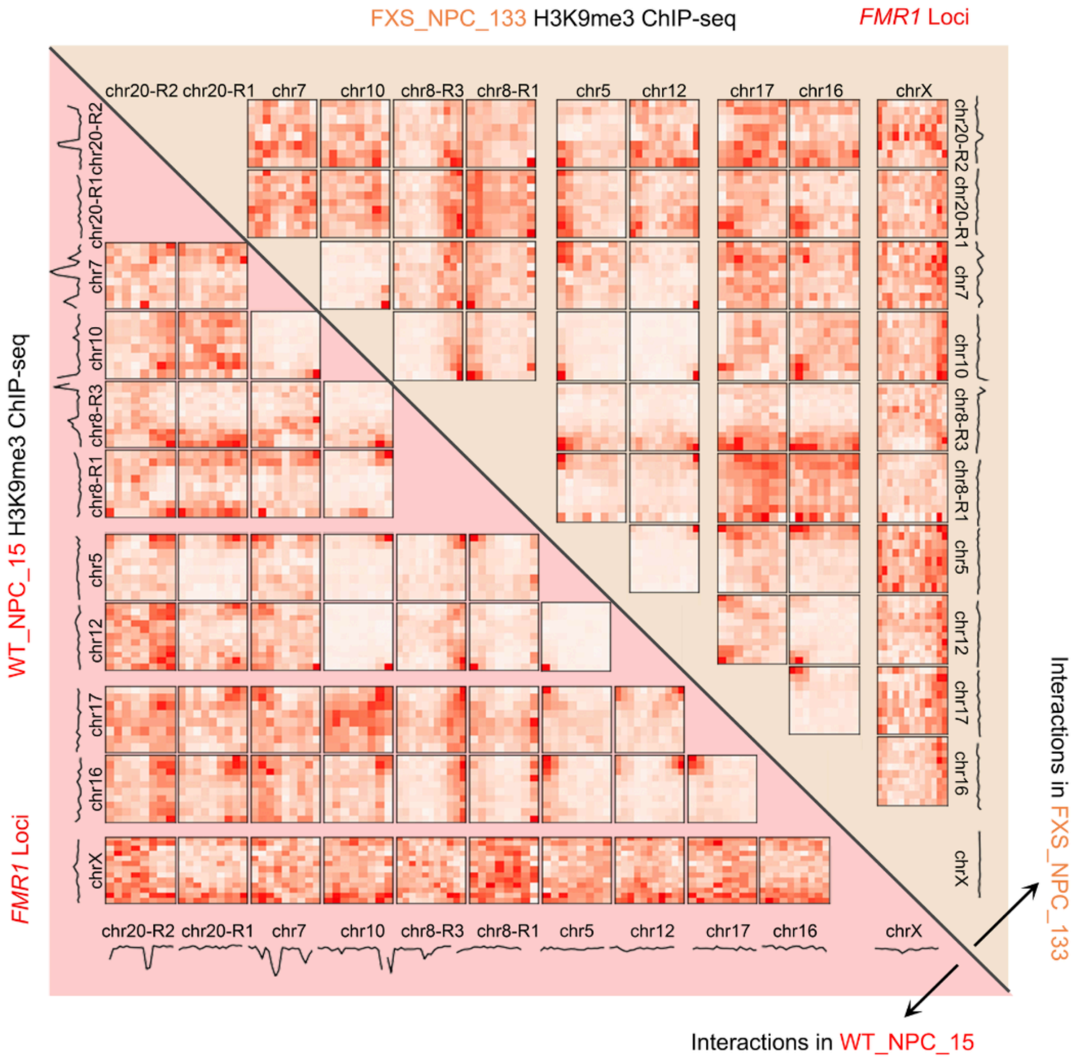


**Fig. S16. Inter-chromosomal interactions between distal H3K9me3 domains in FXS_NPC_15 and FXS_NPC_133.**
Pairwise Hi-C interactions among all of the distal H3K9me3 domains gained in FXS for pre-mutation-length (FXS_133, upper triangle) and normal-length (FXS_15, lower triangle). Domains are annotated by chromosome. The window for each region includes the H3K9me3 domain gained and 3 Mb of flanking genome. H3K9me3 ChIP-seq signal for all domains for both FXS_15 and FXS_133 are plotted above Hi-C heatmaps.

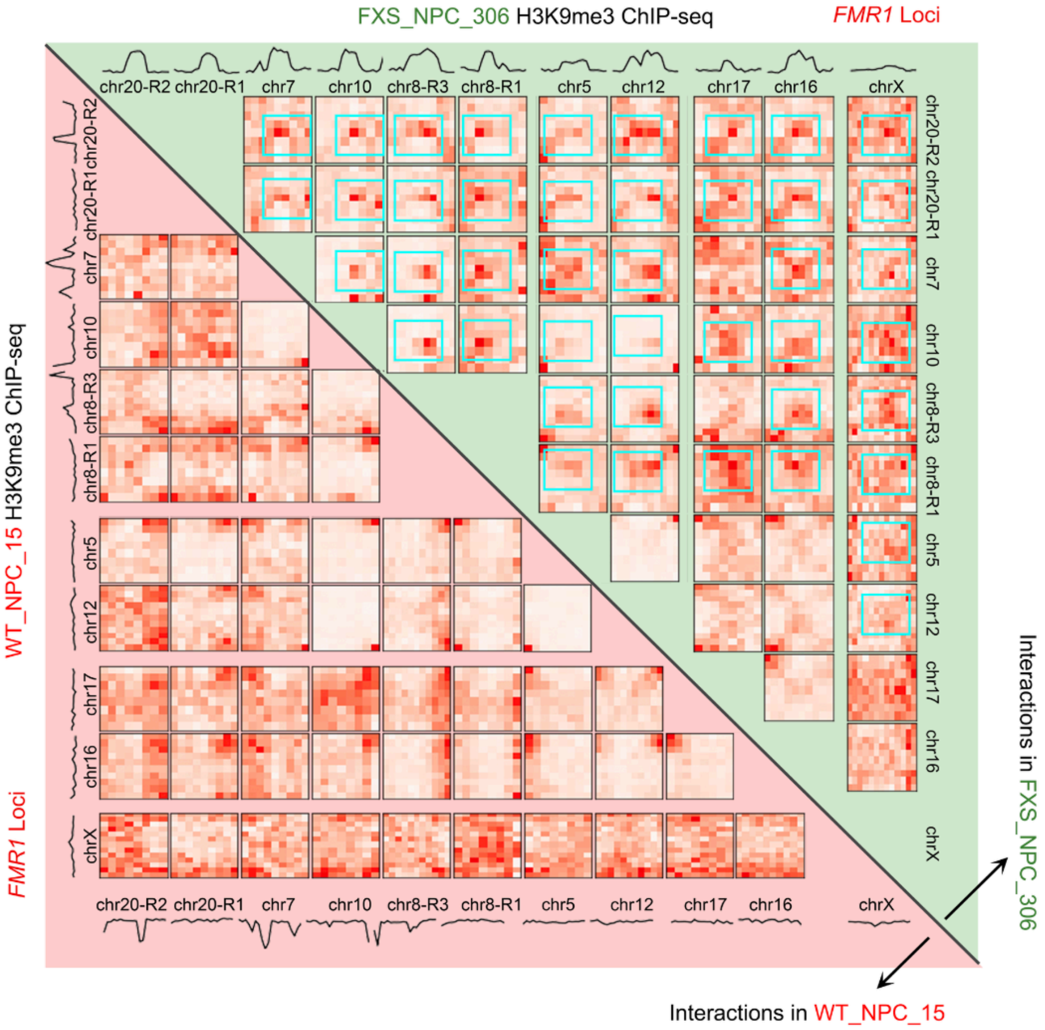


**Fig. S17. Inter-chromosomal interactions between distal H3K9me3 domains in FXS_NPC_15 and FXS_NPC_306.**
Pairwise Hi-C interactions among all of the distal H3K9me3 domains gained in FXS for short mutation-length (FXS_306, upper triangle) and normal length (FXS_15, lower triangle). Domains are annotated by chromosome. The window for each region includes the H3K9me3 domain gained and 3 Mb of flanking genome. H3K9me3 ChIP-seq signal for all domains for both FXS_15 and FXS_306 are plotted above Hi-C heatmaps. Blue boxes highlight gained trans interactions in FXS.

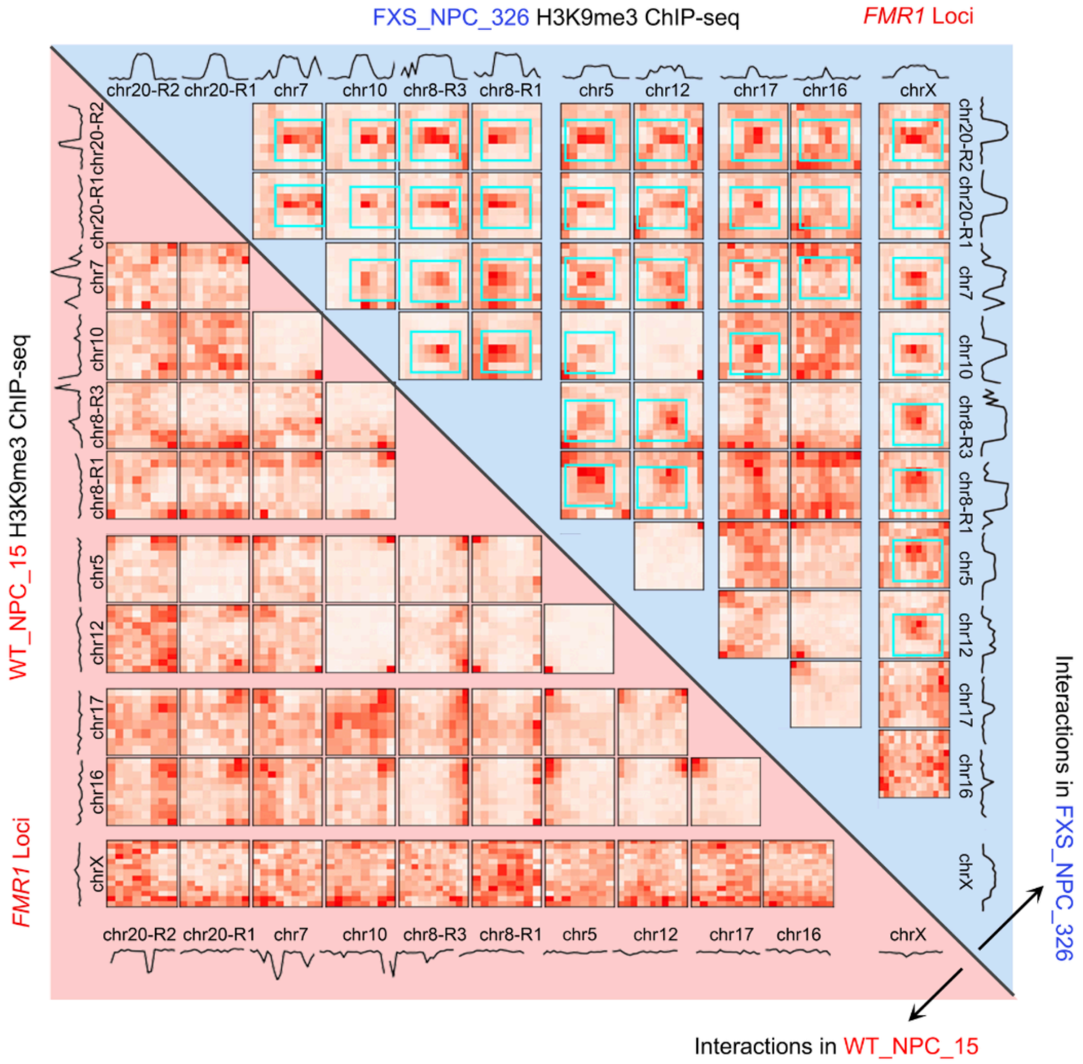


**Fig. S18. Inter-chromosomal interactions between distal H3K9me3 domains in FXS_NPC_15 and FXS_NPC_326.**
Pairwise Hi-C interactions among all of the distal H3K9me3 domains gained in FXS for long mutation-length (FXS_326, upper triangle) and normal length (FXS_15, lower triangle). Domains are annotated by chromosome. The window for each region includes the H3K9me3 domain gained and 3 Mb of flanking genome. H3K9me3 ChIP-seq signal for all domains for both FXS_15 and FXS_326 are plotted above Hi-C heatmaps. Blue boxes highlight gained trans interactions in FXS.

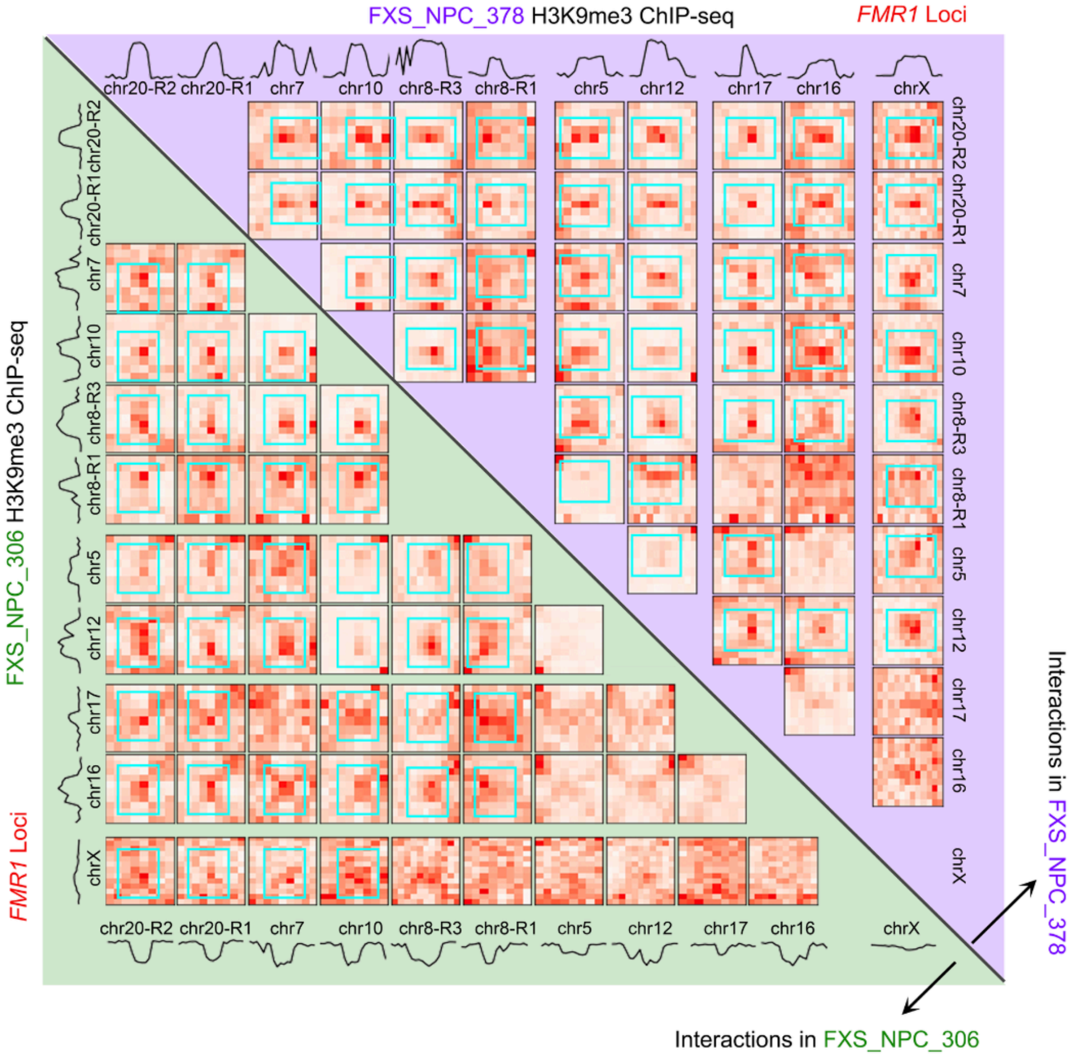


**Fig. S19. Inter-chromosomal interactions between distal H3K9me3 domains in FXS_NPC_306 and FXS_NPC_378.**
Pairwise Hi-C interactions among all of the distal H3K9me3 domains gained in FXS for long mutation-length (FXS_378, upper triangle) and short mutation-length (FXS_306, lower triangle). Domains are annotated by chromosome. The window for each region includes the H3K9me3 domain gained and 3 Mb of flanking genome. H3K9me3 ChIP-seq signal for all domains for both FXS_378 and FXS_306 are plotted above Hi-C heatmaps. Blue boxes highlight gained trans interactions in FXS.

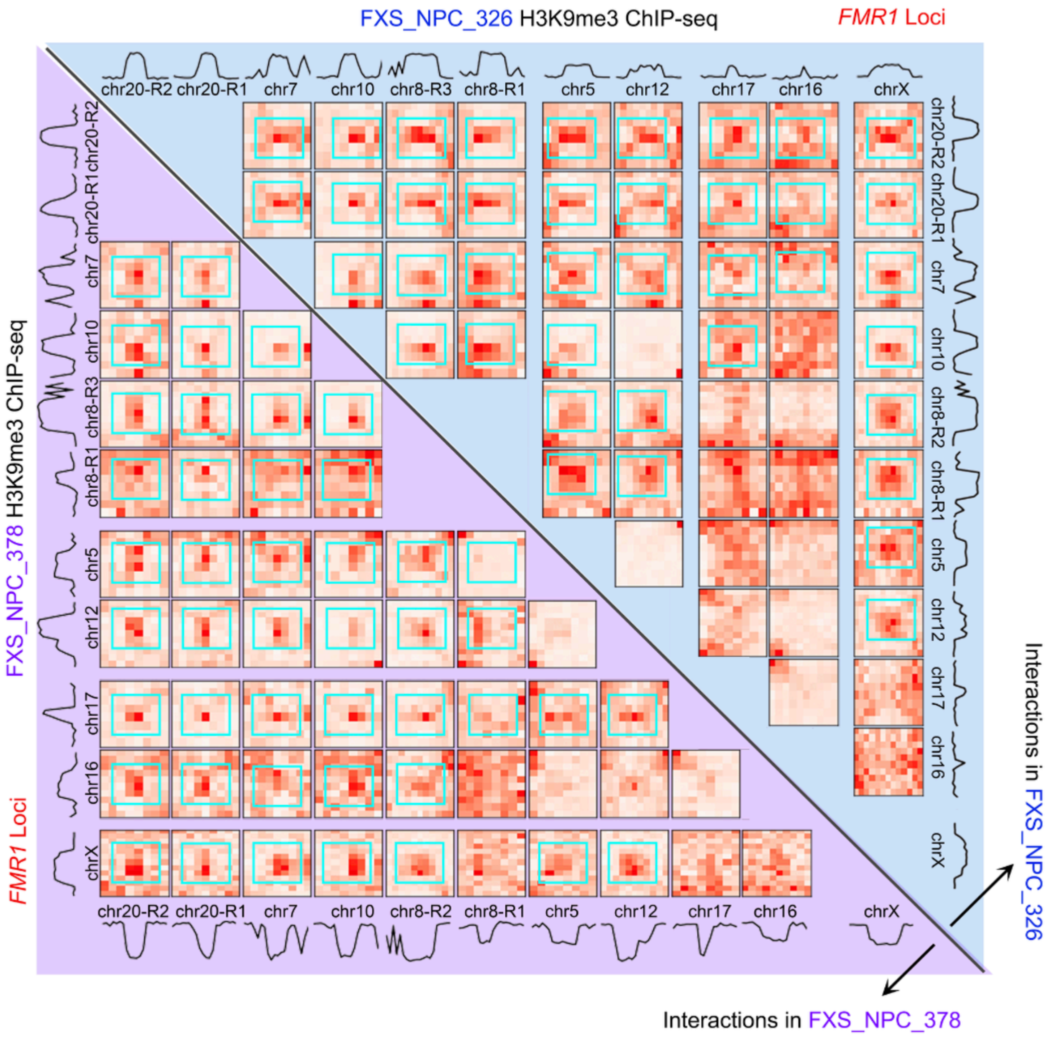


**Fig. S20. Inter-chromosomal interactions between distal H3K9me3 domains in FXS_NPC_326 and FXS_NPC_378.**
Pairwise Hi-C interactions among all of the distal H3K9me3 domains gained in FXS for short mutation-length (FXS_326, upper triangle) and long mutation-length (FXS_378, lower triangle). Domains are annotated by chromosome. The window for each region includes the H3K9me3 domain gained and 3 Mb of flanking genome. H3K9me3 ChIP-seq signal for all domains for both FXS_378 and FXS_326 are plotted above Hi-C heatmaps. Blue boxes highlight gained trans interactions in FXS.

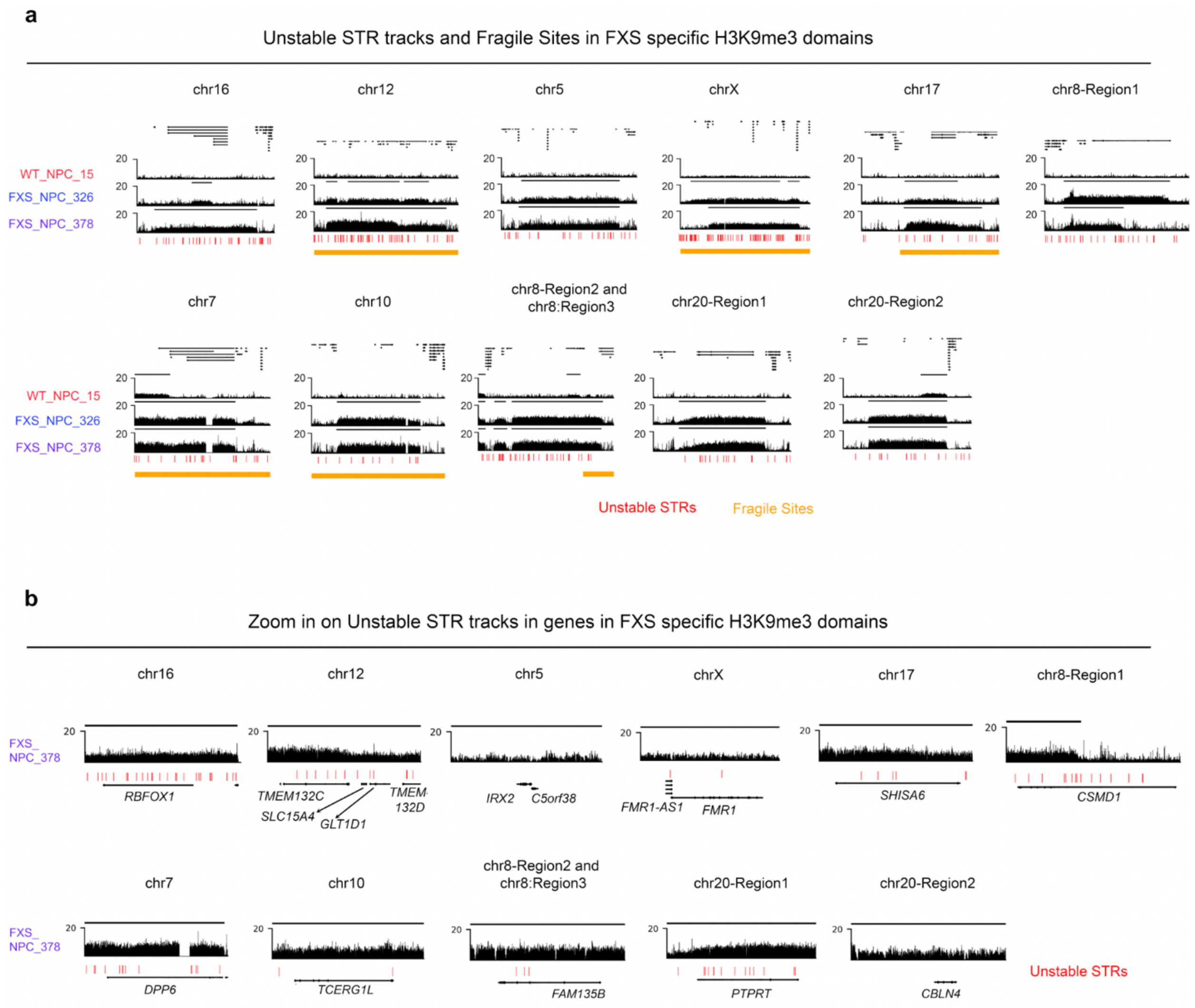


**Fig. S21. Unstable STR tracks and fragile sites with respect to FXS H3K9me3 domains.** (a) For each of FXS specific H3K9me3 domains, the H3K9me3 ChIP-seq in WT_NPC_15, FXS_NPC_326, and FXS_NPC_378 is shown. Underneath, unstable STR tracks are shown in red, fragile sites are shown with orange bars. Fragile sites were obtained from the HumCFS database. (b) Zoom-ins on unstable STR tracks at genes in each of the FXS H3K9me3 domains.

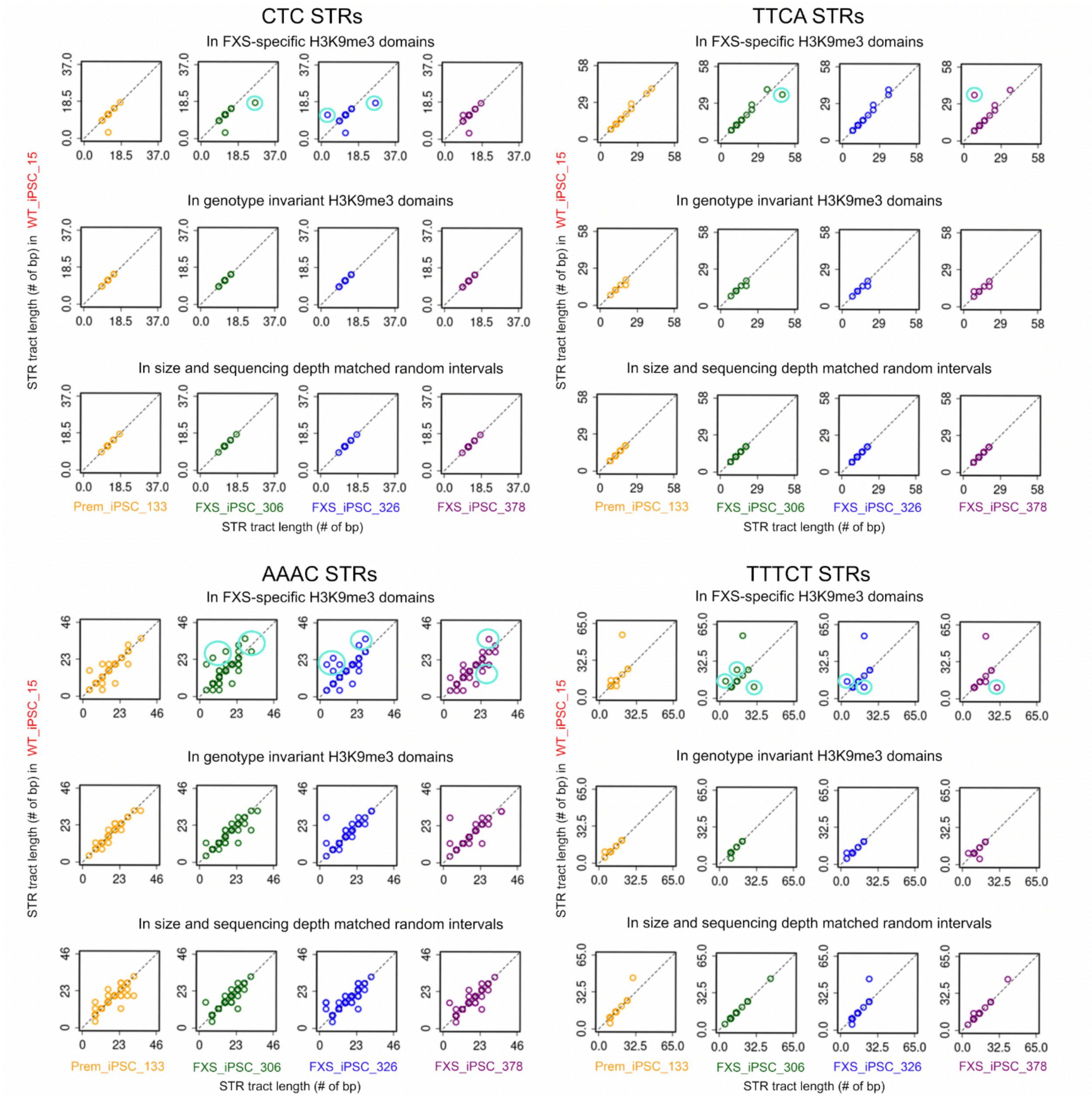


**Fig. S22. STRs which change length in FXS specific domains in FXS cell lines.**
For each STR type listed, the length of all those STR loci within either n=12 FXS-specific H3K9me3 domains, n=12 genotype invariant H3K9me3 domains, and n=12 size and sequencing depth matched random intervals are plotted in WT_iPSC_15 versus premutation or disease cell lines.

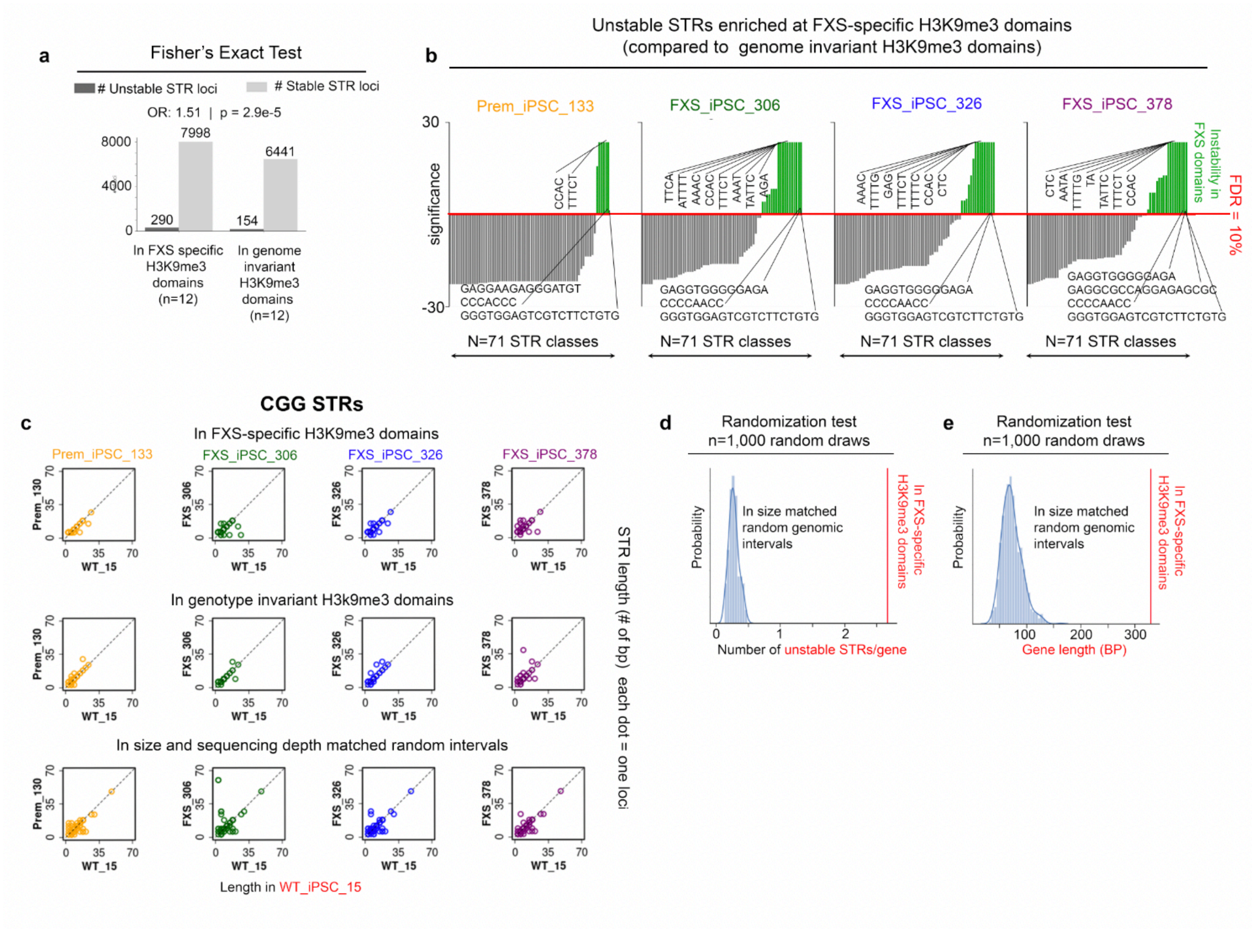


**Fig. S23. Statistical tests demonstrating unique genetic features of FXS H3K9me3 domains compared to the rest of the genome.**

(a) Fisher's Exact test comparing the number of stable vs unstable STR loci in FXS specific H3K9me3 domains vs an equal number coverage matched genome invariant H3K9me3 domains. (b) Results of a statistical test (see Supplemental Methods) demonstrating which of the N=71 classes of STRs which are unstable in FXS are enriched for instability in FXS-specific H3K9me3 domains vs genome invariant H3K9me3 domains. An FDR of 10% was used as a threshold for significance. The individual N=71 loci are plotted on the x-axis from least to most significant. The y-axis, significance, is -10*log(BH-corrected p-value + pseudocount), centered so that FDR of less than 10% is positive and the remaining are negative. (c) For all CGG STRs, the length of all those STR loci within either n=12 FXS-specific H3K9me3 domains, n=12 genotype invariant H3K9me3 domains, and n=12 size and sequencing depth matched random intervals are plotted in WT_iPSC_15 versus premutation or disease cell lines. (d) The number of unstable STRs/gene in n=12 FXS-specific H3K9me3 domains is compared to the number of unstable STRs/gene in n=1,000 random draws of n=12 size matched genomic intervals. (e) The average gene length in BP in n=12 FXS-specific H3K9me3 domains is compared to the average gene length in n=1,000 random draws of n=12 size matched genomic intervals.

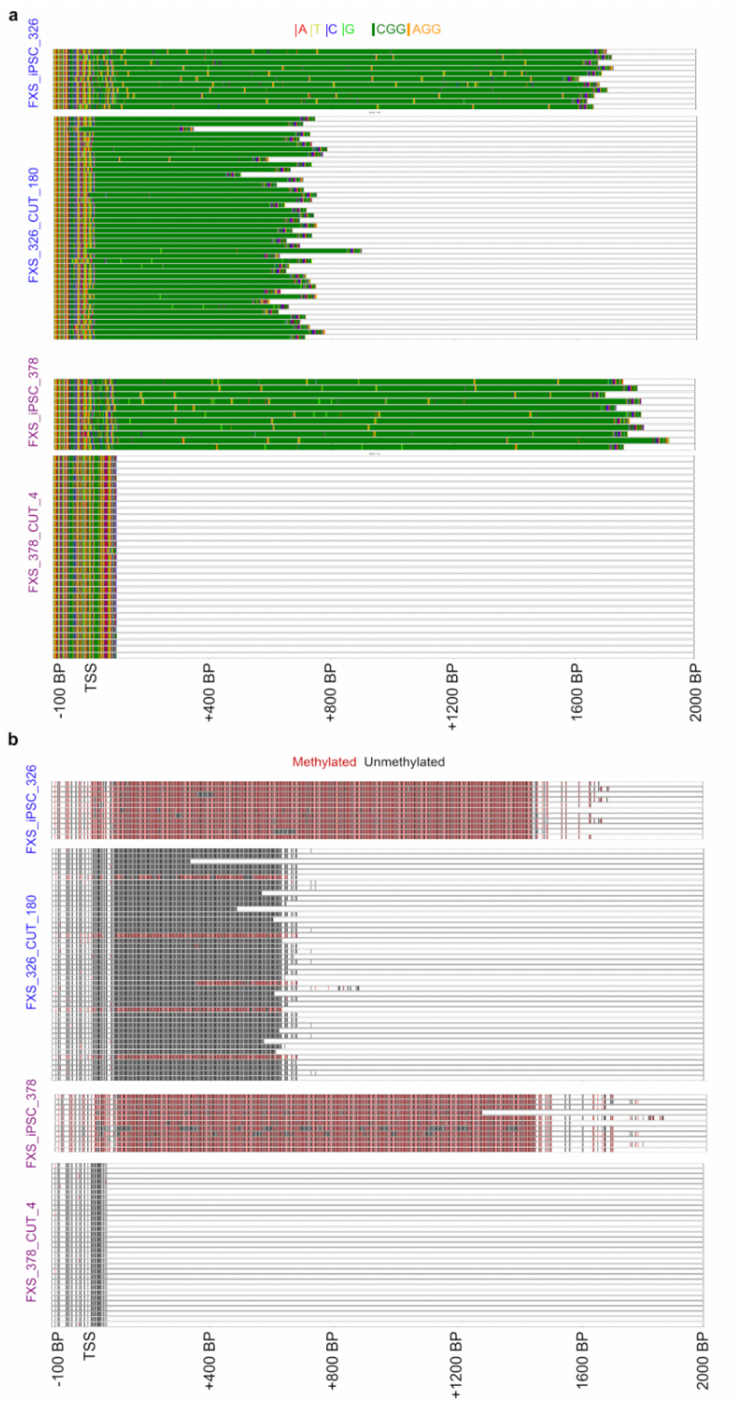


**Fig. S24. Nanopore long-read sequencing and CpG methylation analysis of edited cell lines.** (a) Visual representation of Nanopore long reads that span the transcription start site and first 200 bp of *FMR1*. For each of the 4 samples, the sequence of each read is shown with colors corresponding to base pairs as shown in the legend (top right). (b) For each long read obtained per cell line, DeepSignal was used to determine the methylation status of each CpG present in the sequence (see Methods). Black indicates an unmethylated CpG and red indicates a methylated CpG. Blanks indicate no CpG present.

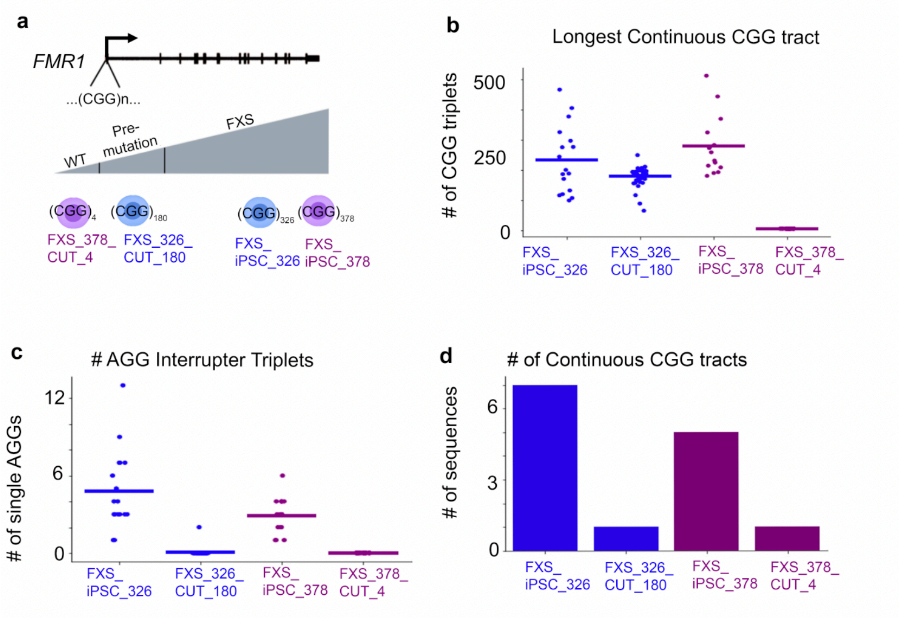


**Fig. S25 Length characterization of CRISPR edited iPSC lines.**
(a) Schematic of iPSC lines and the CRISPR edited deletions used in this study. (b-e) Nanopore long-read analysis of (b) longest continuous CGG tract, (c) number of AGG interrupters within CGG STR, and (d) total number of continuous CGG tracks within the STR in the 5'UTR of *FMR1*.

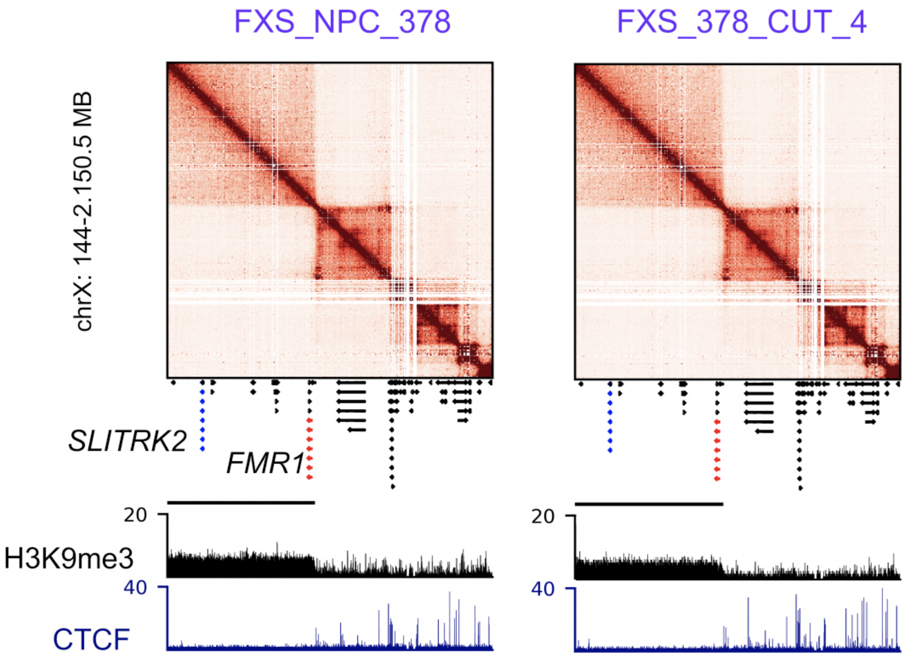


**Fig. S26. 5C, H3K9me3, and CTCF in fragile X syndrome in iPSC upon CGG repeat cut to normal length.**
5C in a ~6 Mb region around *FMR1* is shown for a disease cell line, FXS_iPSC_378, and an isogenic line where the repeats were cut out from disease length to normal length (378_CUT_4). H3K9me3 ChIP-seq and CTCF for each are shown below each heatmap.

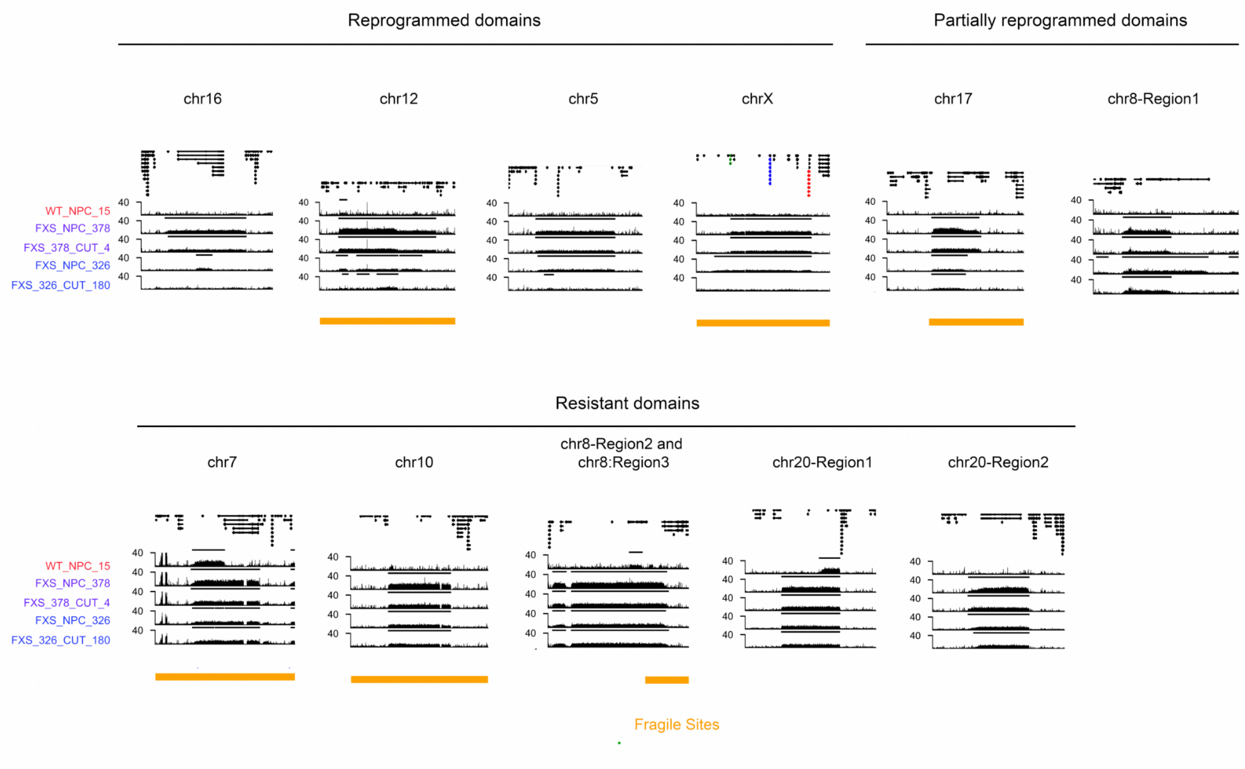


**Fig. S27. Distal H3K9me3 domains in iPSC FXS upon CGG repeat cut out.**
H3K9me3 Chip-seq is shown around distal FXS specific H3K9me3 domains in one replicate for of each cell line in Isogenic set 1 and set 2. Lines representing RSEG H3K9me3 domain calls are shown above H3K9me3 ChIP-seq track. Red represents locations of CGGCGG, and yellow represents fragile sites, obtained from HumFCS database.

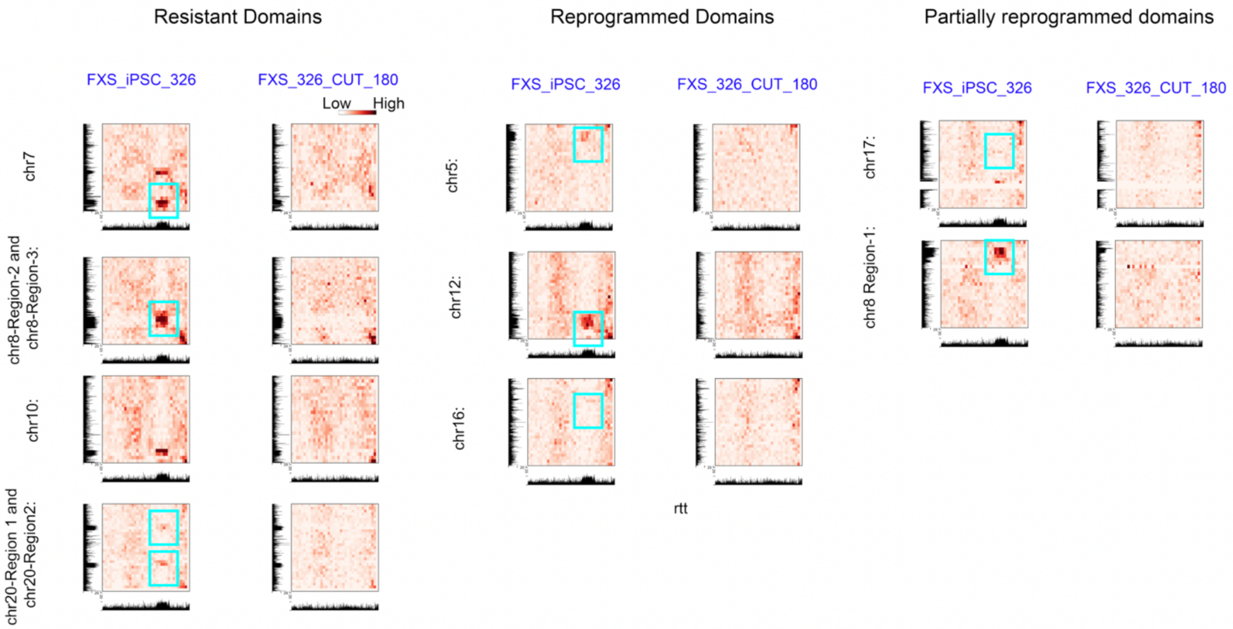


**Fig. S28. Inter-chromosomal interactions between chr X (*FMR1* loci) and distal H3K9me3 domains in FXS_iPSC_326 and isogenic cut out line FXS_326_CUT_180.**
Hi-C interactions between *FMR1* loci on chromosome X and the distal H3K9me3 domains (see Figure 3) are shown for long mutation-sample FXS_iPSC_326 and the edited cell lines with CGGs cut to 180, (FXS_326_CUT_180). The window for each region includes the H3K9me3 domain gained and up to 20 Mb of flanking genome. H3K9me3 ChIP-seq *FMR1* is shown on the x axis (chr X) and for the distal region is shown on the Y axis. All data is from one replicate per cell line. Hi-C data is binned at 1 Mb resolution. The domains are identified by which chromosome they are on, and the *FMR1* domain is identified as chr X.

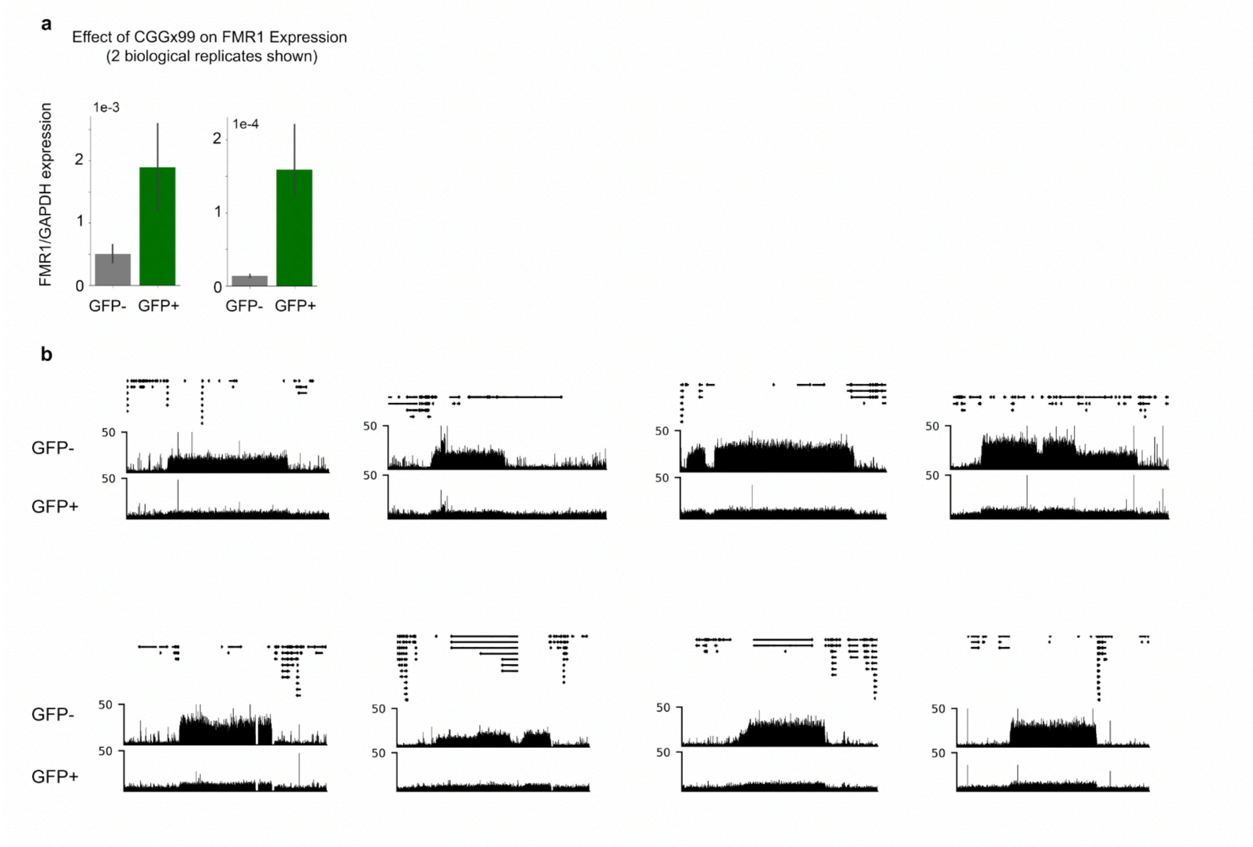


**Fig. S29. Effect of CGG-99x on gene expression and H3K9me3 domain strength.**
(a) Expression of *FMR1* via qRT-PCR in cells which either did receive (GFP+) or did not receive (GFP-) the CGGx99 plasmid. Two biological replicates separate from that in Figure 5b are shown. (b) H3K9kme3 ChIP-seq in consistently gained H3K9me3 domains in FXS.
